# ER Proteostasis Regulators Reduce Amyloidogenic Light Chain Secretion Through an On-Target, ATF6-Independent Mechanism

**DOI:** 10.1101/2020.05.15.098145

**Authors:** Bibiana Rius, Jaleh S. Mesgarzadeh, Isabelle C. Romine, Ryan J. Paxman, Jeffery W. Kelly, R. Luke Wiseman

## Abstract

The plasma cell secretion and toxic aggregation of amyloidogenic immunoglobulin light chains (LCs) causes proteotoxicity in Light Chain Amyloidosis (AL). We recently identified endoplasmic reticulum (ER) proteostasis regulators such as compound **147** that reduce secretion and aggregation of LCs implicated in AL (Plate, Cooley et al., 2016). Compound **147** promotes adaptive ER proteostasis remodeling through a mechanism involving covalent modification of multiple protein disulfide isomerases (PDIs) and subsequent activation of the ATF6 unfolded protein response (UPR) -associated transcriptional signaling pathway (Paxman, Plate et al., 2018). Here, we show that the **147**-dependent reduction in amyloidogenic LC secretion from AL patient plasma cells is independent of ATF6 activation, but instead requires on-target PDI modification. Our results reveal pharmacologic targeting of PDIs as a potential strategy to ameliorate AL-associated proteotoxicity and demonstrate that **147** can influence ER proteostasis through multiple on-target mechanisms including ATF6 activation and PDI modification.

**IMPACT STATEMENT:** This study demonstrates the broad potential for endoplasmic reticulum proteostasis regulator compounds such as **147** to influence secretory proteostasis of disease-associated proteins through multiple on target mechanisms.

## INTRODUCTION

Light chain amyloidosis (AL) is the most common systemic amyloid disease, affecting 8-10 people per million per year (Badar, D’Souza et al., 2018, Blancas-Mejia & Ramirez-Alvarado, 2013, Cohen & Comenzo, 2010, Merlini, Comenzo et al., 2014, Merlini, Dispenzieri et al., 2018). AL pathogenesis involves the toxic extracellular aggregation of an amyloidogenic immunoglobulin light chain (LC) that is secreted from a clonally expanded cancerous plasma cell. These LC aggregates deposit on post-mitotic tissues including the heart and kidneys causing organ failure and ultimately death (Badar et al., 2018, Blancas-Mejia, Misra et al., 2018, Brenner, Jain et al., 2004, Diomede, Romeo et al., 2017, Guan, Mishra et al., 2015, Guan, Mishra et al., 2013, Imperlini, Gnecchi et al., 2017, Merlini et al., 2014, Merlini et al., 2018, Shi, Guan et al., 2010). Current treatments for AL use chemotherapy combined with autologous stem cell replacement to eliminate the AL-associated plasma cells (Badar et al., 2018, Gertz, 2018, Kastritis, Dialoupi et al., 2019, Manwani, Hegenbart et al., 2018, Merlini et al., 2018, Milani, Palladini et al., 2018, Pinney, Smith et al., 2013). This reduces circulating serum populations of amyloidogenic LCs, decreasing toxic LC aggregation, promoting clearance of deposited amyloid, and improving organ function (Dember, Sanchorawala et al., 2001, Meier-Ewert, Sanchorawala et al., 2011, Merlini et al., 2018, Salinaro, Meier-Ewert et al., 2017). However, ~30% of AL patients presenting with severe cardiac or renal LC proteotoxicity are too ill at diagnosis to tolerate chemotherapy (Merlini et al., 2018, Palladini, Hegenbart et al., 2014, Wechalekar, Schonland et al., 2013). Thus, new strategies are required to alleviate the LC proteotoxicity on distal tissues and allow chemotherapeutic access to AL patients with severe organ involvement.

One strategy to reduce the secretion and toxic aggregation of amyloidogenic proteins is through the adaptive remodeling of the ER proteostasis network comprising ER chaperones (e.g., BiP), folding enzymes (e.g., protein disulfide isomerases, PDIs), and degradation factors (Plate & Wiseman, 2017, Powers, Morimoto et al., 2009). These ER proteostasis pathways function to partition ER proteins between folding, trafficking, and degradation in a process termed ER quality control (Braakman & Bulleid, 2011, Plate & Wiseman, 2017, Sun & Brodsky, 2019). Through this partitioning, ER proteostasis pathways reduce the secretion and aggregation of non-native, aggregation-prone proteins in secretory environments including the ER and extracellular space. In the context of AL, destabilized amyloidogenic LCs escape plasma cell ER quality control, allowing their efficient secretion to the serum (Blancas-Mejia et al., 2018, Eisele, Monteiro et al., 2015, Plate & Wiseman, 2017). This increases extracellular populations of amyloidogenic LCs available for concentration-dependent aggregation into the toxic oligomers and amyloid fibrils implicated in AL pathogenesis.

Enhancing ER proteostasis through mechanisms such as activation of the unfolded protein response (UPR)-associated transcription factor ATF6 can selectively reduce the secretion and toxic aggregation of destabilized, amyloidogenic proteins (Kelly, 2020, Plate & Wiseman, 2017). ATF6 induces the expression of many ER proteostasis factors including the ATP-dependent ER HSP70 BiP and multiple protein disulfide isomerases (PDIs) (Haze, Yoshida et al., 1999, Shoulders, Ryno et al., 2013). Genetic activation of ATF6 using a ligand-regulated system preferentially reduces secretion and subsequent aggregation of a destabilized, amyloidogenic LC from HEK293T cells, without impacting the secretion of a non-amyloidogenic LC, fully-assembled IgGs, or the endogenous secretory proteome (Cooley, Ryno et al., 2014, Plate, Rius et al., 2019, Shoulders et al., 2013). Interestingly, the ATF6-dependent reduction in amyloidogenic LC secretion results from increased interactions with ER chaperones and PDIs, which preferentially retain the amyloidogenic LC within the ER and prevent its secretion to downstream secretory environments (Plate et al., 2019). This indicates that pharmacologic approaches that similarly target ER proteostasis could also reduce secretion and toxic aggregation of amyloidogenic LCs implicated in AL (Kelly, 2020, Plate & Wiseman, 2017).

Recently, we used high-throughput screening to identify small molecule ER proteostasis regulators that selectively activate the ATF6 UPR signaling pathway (Plate et al., 2016). One of the most promising small molecules emerging from this screen was compound **147**. Using a combination of medicinal chemistry and chemical biology, we showed that **147** activates ATF6 through a mechanism involving compound metabolic activation and covalent labeling of multiple PDIs (Paxman et al., 2018). Since PDIs are involved in regulating the proper folding and disulfide bond formation for ER-targeted proteins (Appenzeller-Herzog & Ellgaard, 2008, Braakman & Bulleid, 2011, Wang, Wang et al., 2015), this mechanism of action suggested that **147** can influence ER proteostasis through two on-target mechanisms: covalent PDI modification and activation of the UPR-associated transcription factor ATF6. Previous work shows that **147** protected cardiomyocytes both in vitro and in vivo from pathologic insults through an ATF6-dependent mechanism (Blackwood, Azizi et al., 2019), highlighting the potential for **147**-dependent ATF6 activation to promote protective signaling. However, the potential to influence ER proteostasis of secretory proteins through **147**-dependent PDI modification had not been previously demonstrated.

Here, we show that compound **147** reduces the secretion of the destabilized, amyloidogenic λ6a LC ALLC from AL patient-derived plasma cells. However, this **147**-dependent reduction in ALLC secretion is refractory to co-treatments with ATF6 inhibitors, demonstrating that this compound reduces ALLC secretion through an ATF6-independent mechanism. Instead, our results indicate that **147** reduces ALLC plasma cell secretion through an on-target mechanism involving metabolic activation and covalent modification of multiple PDIs. Consistent with this, other PDI inhibitors that covalently modify a similar subset of PDIs also reduce ALLC plasma cell secretion through a mechanism analogous to that observed for **147**. These results indicate that pharmacologic targeting of PDIs using ER proteostasis regulators like **147** represents a potential strategy to mitigate the plasma cell secretion of amyloidogenic LCs implicated in AL. Importantly, we further show that **147** is compatible with chemotherapeutics used to ablate AL plasma cells (e.g., bortezomib), indicating that these two approaches could be used in combination to reduce both the LC-associated organ toxicity and underlying plasma cell malignancy observed in AL pathogenesis. Ultimately, our results demonstrate that, apart from promoting ATF6-mediated ER proteostasis remodeling, compound **147** can also correct pathologic imbalances in ER proteostasis through the covalent targeting of PDIs. This highlights the broad potential for this compound to influence ER proteostasis for structurally diverse proteins implicated in many types of protein misfolding disease.

## RESULTS

### Compound 147 reduces ALLC secretion from AL patient plasma cells

We previously showed that **147** reduces secretion of the amyloidogenic λ6a LC ALLC from AL patient-derived ALMC-2 cells by 50% (Plate et al., 2016). We confirmed this **147**-dependent reduction in ALLC secretion by both immunoblotting and ELISA (**Figure 1A** and **Figure 1 – figure supplement 1A**). However, treatment with **147** did not significantly influence the viability of ALMC-2 cells, indicating that this reduced secretion cannot be attributed to cell death (**Figure 1 – figure supplement 1A**). Compound **147** also did not significantly reduce IgG secretion or cell viability in Multiple Myeloma (MM)-derived KAS-6/1 cells (**Figure 1 – figure supplement 1B**). This supports previous findings showing that this compound preferentially reduces secretion of destabilized, amyloidogenic LCs (Plate et al., 2016). Amyloidogenic λ LCs are secreted in multiple conformations including monomers, disulfide-bound dimers, and fully assembled IgGs (Kaplan, Ramirez-Alvarado et al., 2009). However, the monomers and dimers are the predominant species associated with toxic LC aggregation and AL amyloid pathology (Andrich, Hegenbart et al., 2017, Brenner et al., 2004, Klimtchuk, Gursky et al., 2010, Ramirez-Alvarado, 2012, Sikkink & Ramirez-Alvarado, 2010). Using non-reducing SDS-PAGE, we showed that **147** reduces secretion of ALLC monomers and dimers from ALMC-2 cells (**Figure 1B,C**). This demonstrates that this compound decreases extracellular populations of the species most associated with toxic LC aggregation. Using a cycloheximide (CHX) chase assay, we showed that **147** reduces the fraction of ALLC secreted by 30% (**Figure 1D**). However, this reduction in ALLC secretion did not correspond to a significant reduction in total ALLC over the 6 h time course of this experiment, indicating that **147** does not significantly increase ALLC degradation (**Figure 1E**). This result is consistent with previous results, demonstrating that both stress-independent ATF6 activation and treatment with **147** reduces ALLC secretion through a mechanism involving its increased intracellular retention in complexes bound to ER chaperones such as BiP (Plate et al., 2016, Plate et al., 2019).

**Figure 1 (with 1 supplement).**
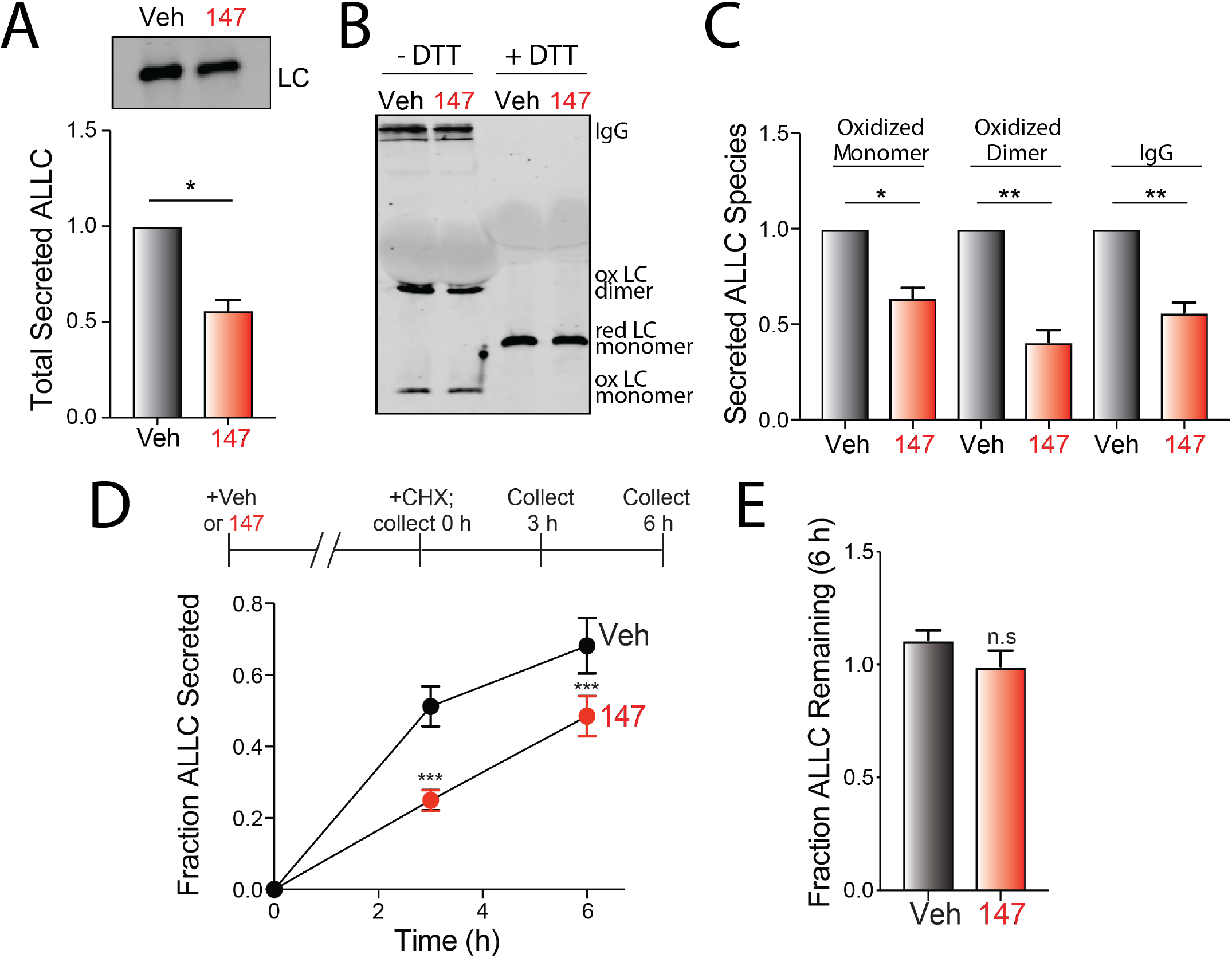
Compound 147 reduces secretion of ALLC from AL patient derived ALMC-2 cells. **A.** Representative immunoblot and quantification of ALLC in conditioned media prepared on ALMC-2 cells treated for 18 hr with vehicle or **147** (10 μM). Error bars show SEM for n= 4 independent experiments. *p<0.05 for a paired t-test. **B.** Representative non-reducing (−DTT) and reducing (+DTT) immunoblots of conditioned media prepared on ALMC-2 cells treated for 18 hr with vehicle or **147** (10 μM). The fully-assembled IgGs, oxidized LC dimers, oxidized LC monomers, and reduced LC monomers are indicated. **C.** Quantification of non-reducing immunoblots as shown in **Figure 1B** showing the relative recovery of oxidized LC monomers, oxidized LC dimers, and fully-assembled IgGs. Error bars show SEM for n=3 or 4 independent experiments. **p<0.05, **p<0.01 from a paired t-test. **D.** Graph showing the fraction ALLC secreted from ALMC-2 cells treated with vehicle or **147** (10 μM) for 18 hr and then treated with cycloheximide (CHX; 50 μg/mL) at 0, 3 or 6 h. ALLC in conditioned media and lysates were measured by ELISA. The experimental protocol is shown above. Fraction secreted was calculated using: [fraction secreted = ALLC in media at t= 3 or 6 h / ALLC in the lysate at t=0 h]. Error bars show SEM for n=5 replicates. ***p<0.005 vs Veh from an unpaired t-test. **E.** Graph showing the fraction ALLC remaining from ALMC-2 cells treated for 18 h with vehicle or **147** (10 μM) and then treated with cycloheximide (CHX; 50 μg/mL), as in **Figure 1D**. ALLC in conditioned media and lysates were measured by ELISA. Fraction ALLC remaining was calculated using: [fraction ALLC remaining = (ALLC in media t=6 h + ALLC in lysate at t=6h) / ALLC in lysate at t = 0 h]. Error bars show SEM for n= 5 replicates. (n.s), not significant.

### 147 reduces intracellular ALLC in ALMC-2 cells

Although **147** did not significantly increase ALLC degradation in our CHX experiment, we observed 30-50% reductions in intracellular ALLC in **147**-treated ALMC-2 cells by ELISA and immunoblotting (**Figure 2A** and **Figure 2 – figure supplement 1A**). Co-treatment of **147** with either the proteasome inhibitor MG132 or the lysosome inhibitor chloroquine for 5 h did not influence reductions in intracellular or secreted ALLC (**Figure 2 – figure supplement 1A,B**), further indicating that **147** does not increase ALLC degradation over this time course (**Figure 1E**) (Plate et al., 2016). We also did not observe increased recovery of ALLC in cell pellets prepared from **147**-treated ALMC-2 cells, indicating that **147** does not promote intracellular ALLC aggregation (**Figure 2 – figure supplement 1C**). Further, compound **147** did not alter *ALLC* mRNA levels, indicating that reductions in intracellular ALLC cannot be attributed to reduced transcription (**Figure 2 – figure supplement 1D**). However, we did observe a 20% reduction in both newly-synthesized ALLC and total protein in **147**-treated ALMC-2 cells, as measured by [^35^S] metabolic labeling (**Figure 2B-D**). This indicates that the reduced intracellular levels of ALLC can in part be attributed to a decrease in translation. Previous work shows that **147** does not reduce translation in other cells including liver-derived HepG2 cells (Plate et al., 2016), indicating that this reduction in protein synthesis is not universally observed in mammalian cells. Co-treatment with ISRIB, a compound that blocks translational attenuation induced by PERK-eIF2α signaling (Sidrauski, Acosta-Alvear et al., 2013, Sidrauski, McGeachy et al., 2015), does not impact the reduction in secreted or intracellular ALLC in **147**-treated ALMC-2 cells (**Figure 2E,F**). This suggests that the reductions in intracellular ALLC are likely independent of PERK-dependent translation attenuation. Collectively, these results show that **147** reduces intracellular ALLC through a mechanism involving reduced translation and that this reduction in synthesis contributes to the reduced ALLC secreted from compound-treated ALMC-2 cells.

**Figure 2 (with 1 supplement).**
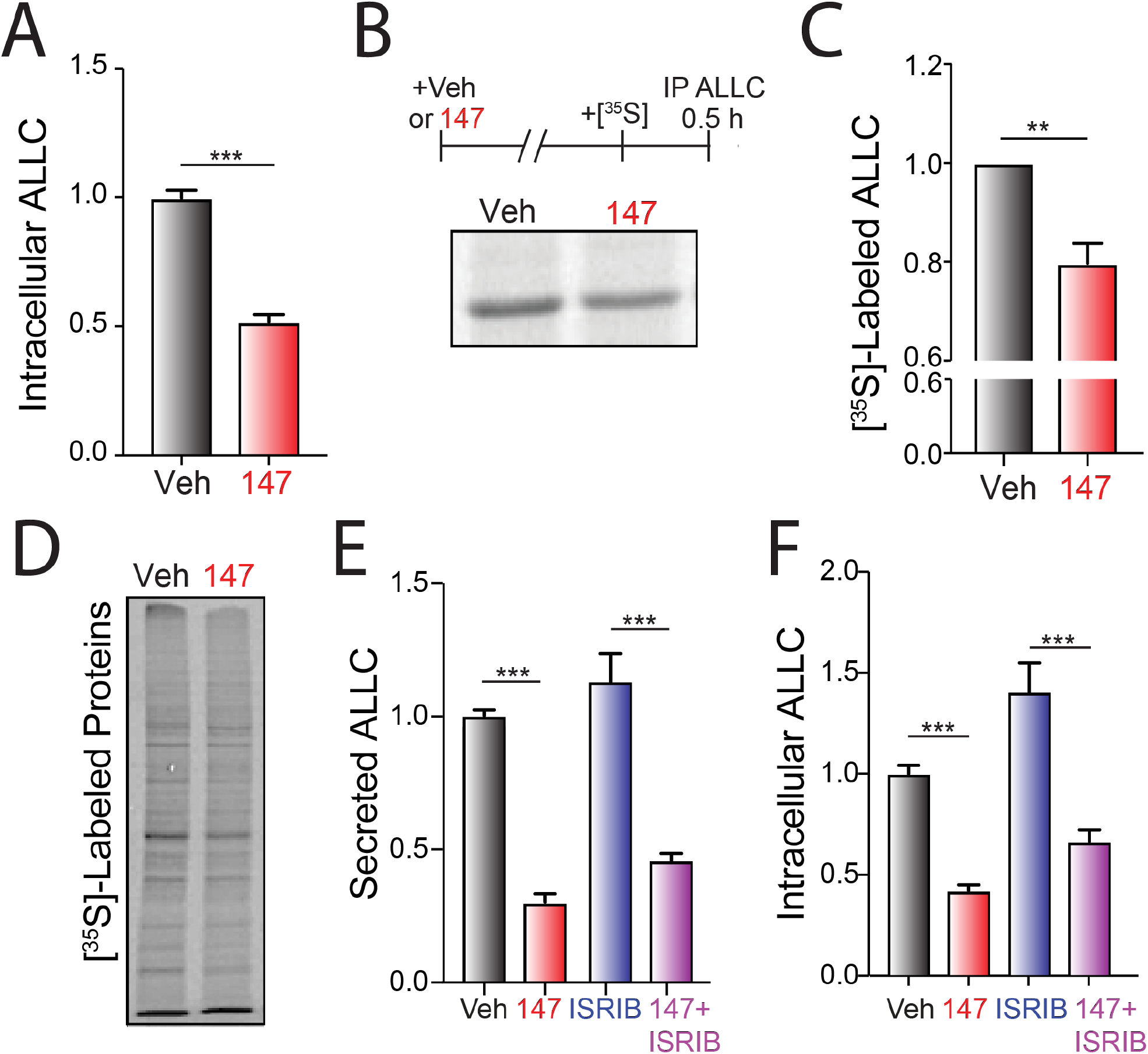
Compound 147 reduces lysate levels of ALLC in ALMC-2 cells. **A.** Graph showing normalized amounts of ALLC in lysates prepared from ALMC-2 cells incubated for 18 hr with vehicle or **147** (10 μM). ALLC was quantified by ELISA. Error bars show SEM for n= 23 replicates across five independent experiments. ***p<0.005 from an unpaired t-test. **B.** Representative autoradiogram of [^35^S]-labeled ALLC immunopurified from ALMC-2 cells treated for 18 hr with vehicle or **147** (10 μM) and then labeled for 30 min with [^35^S]. The experimental protocol is shown above. **C.** ALLC quantification of autoradiograms shown in **Figure 2B** normalized to Veh-treated cells. Error bars show SEM for n= 5 independent experiments. **p<0.01 from a paired t-test. **D.** Autoradiogram of whole cell lysates prepared from ALMC-2 cells treated for 18 hr with vehicle or **147** and then metabolically labeled with [^35^S] for 30 min. **E.** Quantification of ALLC in conditioned media prepared on ALMC-2 cells treated for 18 hr with vehicle, **147** (10 μM), and/or ISRIB (200 nM), as indicated. ALLC was measured by ELISA and normalized to vehicle-treated cells. Error bars show SEM for n = 14 replicates across three independent experiments. ***p<0.005 from an unpaired t-test. **F.** Quantification of ALLC in lysates prepared from ALMC-2 cells treated for 18 hr with vehicle, **147** (10 μM), and/or ISRIB (200 nM), as indicated. ALLC was measured by ELISA and normalized to vehicle-treated cells. Error bars show SEM for n = 14 replicates across three independent experiments. ***p<0.005 from an unpaired t-test.

### 147-dependent reductions in ALLC secretion are not dependent on ATF6 activation

We next defined whether the **147**-dependent reductions in ALLC secretion rely on ATF6 activation. ATF6 is activated through a mechanism involving the reduction of disulfide-linked ATF6 oligomers and increased trafficking of reduced ATF6 monomers to the Golgi (**Figure 3A**) (Koba, Jin et al., 2020, Nadanaka, Okada et al., 2007, Nadanaka, Yoshida et al., 2006, Oka, van Lith et al., 2019). In the Golgi, ATF6 is site-specifically processed by site 1 protease (S1P) and site 2 protease (S2P), releasing the active ATF6 N-terminal bZIP transcription factor domain, which localizes to the nucleus and promotes ATF6 transcriptional activity (Ye, Rawson et al., 2000). Recently, the compound Ceapin-7 (CP7) was shown to stabilize ATF6 oligomers within the ER, preventing ATF6 activation induced by ER stress or ATF6 activating compounds such as **147** (**Figure 3A**) (Gallagher, Garri et al., 2016, Gallagher & Walter, 2016, Paxman et al., 2018, Torres, Gallagher et al., 2019). We confirmed that co-treatment of ALMC-2 cells with CP7, but not the inactive analog CP5 (**Figure 3 – figure supplement 1**), blocked **147**-dependent increases in the ATF6 target gene *BiP* (**Figure 3B**). Interestingly, co-treatment with CP7 did not block **147**-dependent reductions in ALLC secretion from ALMC-2 cells (**Figure 3C**). Further, co-treatment with CP7 did not influence **147**-dependent reductions in intracellular ALLC (**Figure 3D**). This suggests that **147** reduces ALLC secretion through a mechanism independent of ATF6. To further probe this prediction, we co-treated ALMC-2 cells with **147** and the S1P inhibitor PF429242 (S1Pi), a compound that blocks ATF6 activation through a mechanism distinct from CP7 (**Figure 3A**) (Hay, Abrams et al., 2007, Lebeau, Byun et al., 2018). Co-treatment with S1Pi also did not block **147**-dependent reductions in secreted ALLC (**Figure 3E**). These results further confirm that **147** reduces ALLC secretion from ALMC-2 cells through a mechanism independent of ATF6 activation.

**Figure 3 (with 1 supplement).**
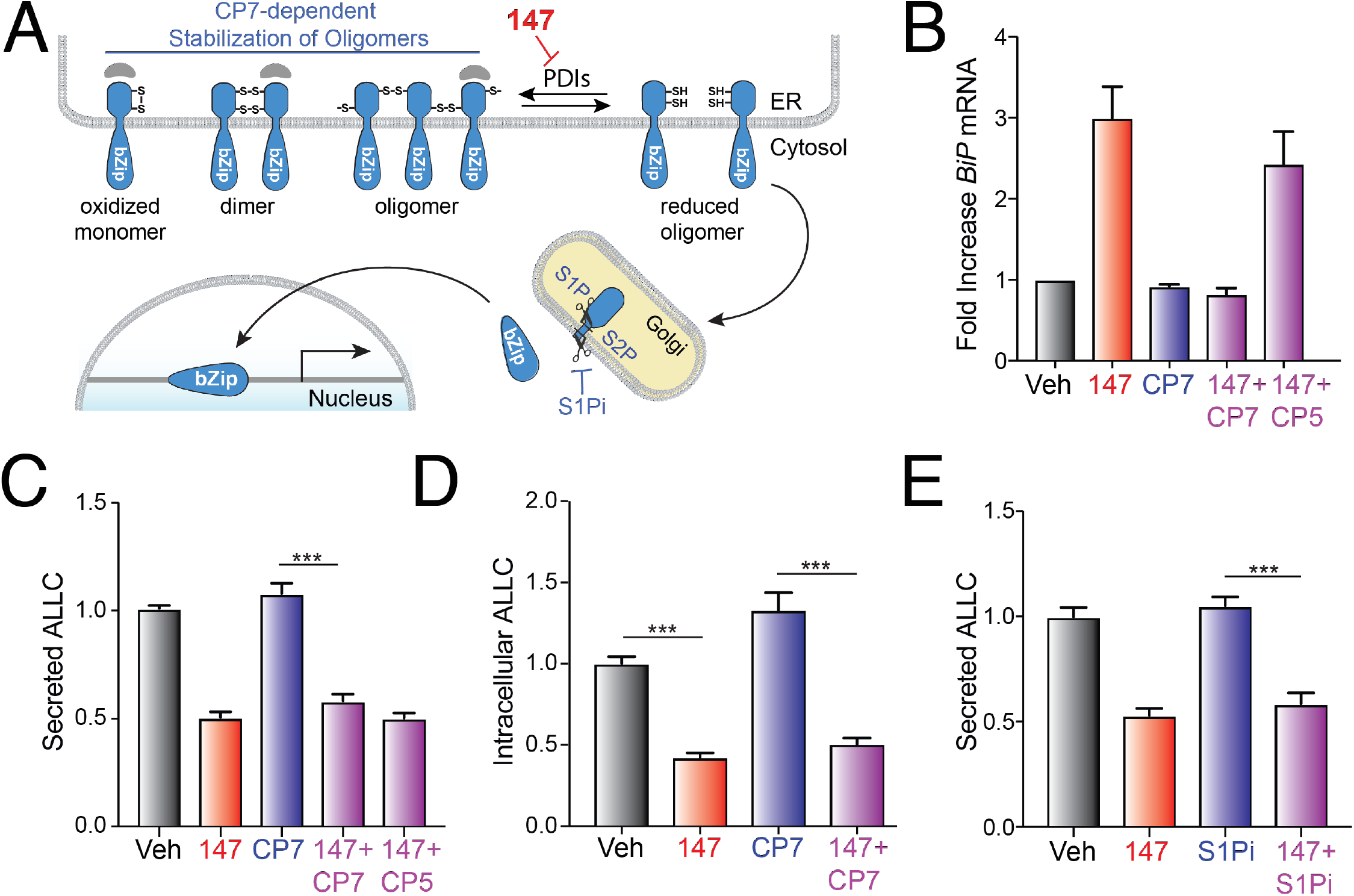
147-dependent reductions in ALLC secretion are independent of ATF6 activation. **A.** Illustration showing the mechanism of **147**-dependent ATF6 activation. ATF6 is retained within the ER as oxidized monomers, disulfide-bonded dimers, or disulfide-bonded oligomers that are maintained by PDIs. Compound **147** covalently modifies a subset of PDIs, increasing the population of reduced ATF6 monomers that can traffic to the Golgi and undergo proteolytic processing by S1P and S2P. This releases the N-terminal bZIP domain of ATF6 to localize to the nucleus and promote ATF6 transcriptional signaling. The specific steps of this activation mechanism inhibited by CP7 or S1Pi are shown. **B.** Graph showing *BiP* expression measured by qPCR in ALMC-2 cells treated for 5 hr with vehicle or **147** (10 μM), CP7 (10 μM), or CP5 (10 μM), as indicated. Error bars show 95% confidence interval for n = 3 replicates. **C.** Bar graphs showing normalized amounts of ALLC in conditioned media prepared from ALMC-2 cells treated for 18 hr with vehicle or **147** (10 μM), CP7 (10 μM), or CP5 (10 μM), as indicated. ALLC was quantified by ELISA. Error bars show SEM for n=21 replicates across four independent experiments. ***p<0.005 from an unpaired t-test. **D.** Bar graphs showing normalized amounts of ALLC in in lysates prepared from ALMC-2 cells treated for 18 hr with vehicle, **147** (10 μM) and/or CP7 (10 μM), as indicated. ALLC was quantified by ELISA. Error bars show SEM for n=10 replicates across two independent experiments. ***p<0.005 from an unpaired t-test. **E.** Bar graphs showing normalized amounts of ALLC in conditioned media from ALMC-2 cells treated for 18 hr with vehicle or **147** (10 μM) and/or the site 1 protease inhibitor PF429242 (S1Pi; 10 μM), as indicated. ALLC was quantified by ELISA. Error bars show SEM for n=8 replicates across two independent experiments. ***p<0.005 from an unpaired t-test.

### 147 reduces ALLC secretion through an on-target mechanism involving metabolic activation and covalent protein modification

Compound **147** activates ATF6 through a mechanism involving metabolic oxidation to a quinone methide (**147-QM**) and subsequent covalent modification of ER-localized proteins including multiple PDIs (**Figure 4A**) (Paxman et al., 2018). This activation mechanism requires the 2-amino-*p*-methyl cresol moiety of the **147** ‘A-ring’ (**Figure 4A**). To define a structure-activity relationship for **147**-dependent reductions in ALLC secretion, we screened **147** analogs to define their ability to reduce ALLC secretion from ALMC-2 cells (Paxman et al., 2018). Compounds containing alterations to the **147** 2-amino-*p*-methyl cresol ‘A-ring’ did not reduce ALLC secretion, indicating that this ring is required for compound activity in this assay (**Figure 4B**). Similarly, analogs that include alterations to the **147** ‘linker’ did not substantially reduce ALLC secretion (**Figure 4A** & **Figure 4 – figure supplement 1A**). However, analogs containing alterations to the **147** ‘B-ring’ showed varying levels of efficacy in reducing ALLC secretion from ALMC-2 cells (**Figure 4A** & **Figure 4 – figure supplement 1B**). This structure-activity relationship is identical to that previously established for **147**-dependent ATF6 activation (Paxman et al., 2018). Consistent with this, we observe an inverse correlation between analog-dependent activation of an ATF6 reporter in HEK293 cells (Paxman et al., 2018) and reductions in ALLC secretion from ALMC-2 cells (**Figure 4C**). This suggests that **147** reduces ALLC secretion from ALMC-2 cells through the same mechanism required for ATF6 activation (**Figure 4A**).

**Figure 4 (with 1 supplement).**
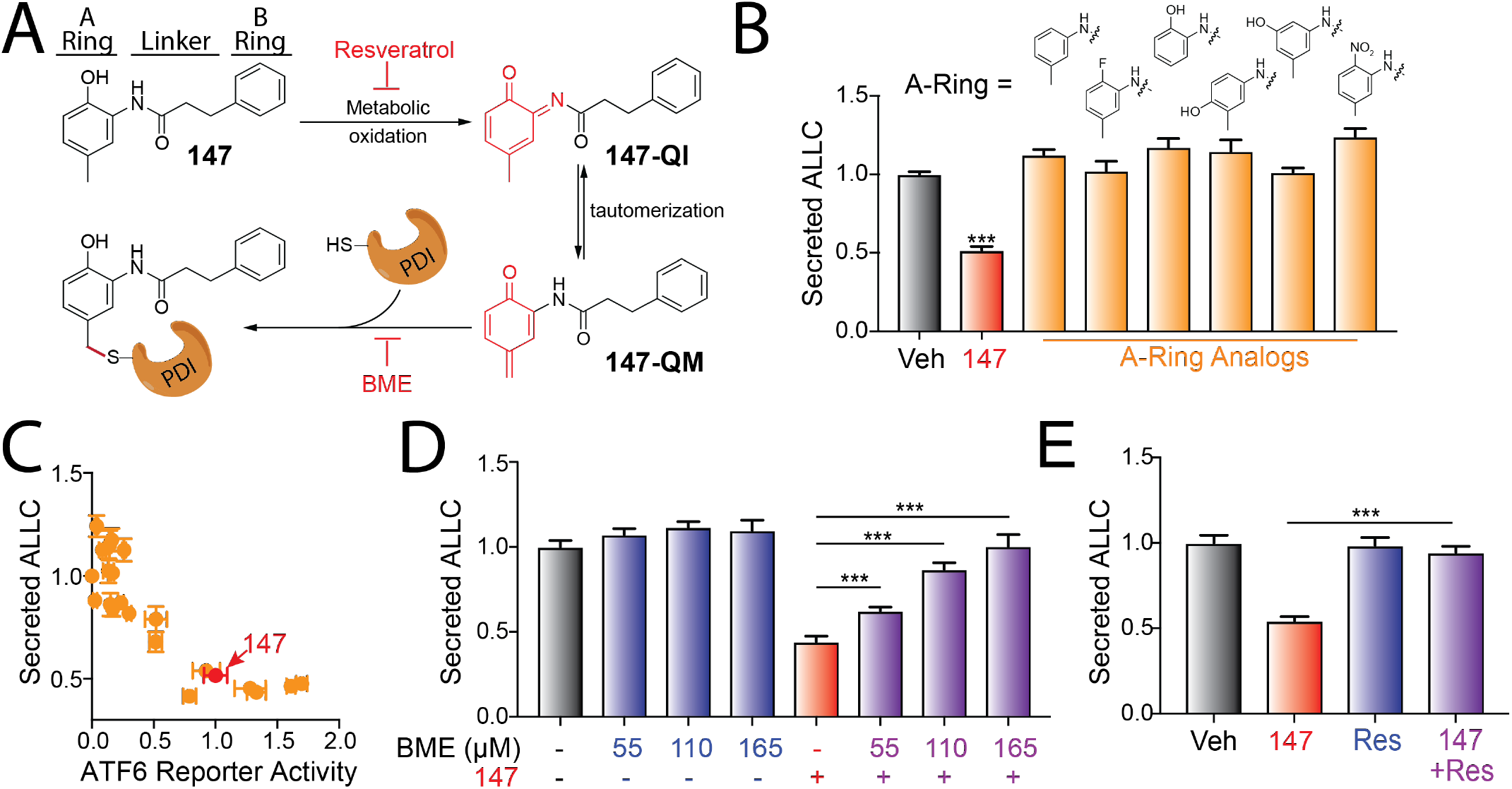
Compound 147 reduces ALLC secretion through a mechanism involving compound metabolic activation and covalent protein modification. **A.** Illustration showing the metabolic activation of **147** to a quinone imide (**147-QI**) and quinone methide (**147**-**QM**). These reactive electrophiles covalently modify ER proteins including multiple PDIs. The ‘A’-ring, linker, and ‘B’-ring of **147** are indicated. Specific steps inhibited by resveratrol and β-mercaptoethanol (BME) are also indicated. Adapted from (Paxman et al., 2018). **B.** Graph showing normalized amounts of ALLC in conditioned media from ALMC-2 cells treated for 18 hr with vehicle, **147** (10 μM) or the indicated **147** ‘A’-ring analog (10 μM). ALLC was quantified by ELISA. Error bars show SEM for n=10 replicates across two independent experiments. ***p<0.005 vs Veh from an unpaired t-test. **C.** Plot comparing the normalized activation of an ATF6-selective luciferase reporter in HEK293 cells (Paxman et al., 2018) to reductions in ALLC secretion from ALMC-2 cells for **147** (10 μM) or **147** analogs containing alterations to the ‘A’-ring, linker, and ‘B’-ring. Error bars for the ATF6 reporter activation show SEM for n=3 independent experiments. Error bars for ALLC secretion show n=10 replicates across two independent experiments. **D.** Graph showing normalized amounts of ALLC secretion from ALMC-2 cells treated for 18 hr with vehicle or **147** (10 μM) and/or β mercaptethanol (BME), as indicated. ALLC was quantified by ELISA. Error bars show SEM for n=10 replicates across two independent experiments. *** p<0.005 from a paired t-test. **E.** Graph showing normalized ALLC secretion from ALMC-2 cells treated for 18 hr with vehicle, **147** (10 μM) and/or resveratrol (Res, 10 μM), as indicated. ALLC was quantified by ELISA. Error bars show SEM for n=9 replicates across two independent experiments. *** p<0.005 from a paired t-test.

The activation of ATF6 by **147** can be inhibited by co-administration of resveratrol or β-mercaptoethanol (BME), which block different steps of the compound activation mechanism (**Figure 4A**) (Paxman et al., 2018). Thus, we sought to define how treatment with resveratrol or BME influences **147**-dependent reduction in ALLC secretion from ALMC-2 cells. Interestingly, co-treatment with BME or resveratrol blocked **147**-dependent reductions in ALLC secretion (**Figure 4D,E**). Neither BME nor resveratrol impacted ALMC-2 cell viability in the absence or presence of **147** (**Figure 4 – figure supplement 1C,D**). Similar results were observed for ALMC-2 cells treated with other active analogs of **147** (**Figure 4 – figure supplement 1E**). Importantly, co-treatment with resveratrol or BME also blocked the **147**-dependent reductions in intracellular ALLC measured by ELISA (**Figure 4 – figure supplement 1F**). Resveratrol also inhibited the translation attenuation in **147**-treated ALMC-2 cells measured by [^35^S] metabolic labeling (**Figure 4 – figure supplement 1G**). These results further demonstrate that **147** reduces ALLC secretion from ALMC-2 cells through the same mechanism required for ATF6 activation, involving both compound metabolic activation and covalent protein modification (**Figure 4A**).

### Compound 147 alters interactions between ALLC and ER PDIs

The above results indicate that **147**-dependent reduction in ALLC secretion from ALMC-2 cells requires covalent modification of proteins. We previously identified proteins modified by **147** using an alkyne-modified analog (**147-alkyne**; **Figure 5 – figure supplement 1A,B**) (Paxman et al., 2018). **147-alkyne** reduces ALLC secretion from ALMC-2 cells to levels identical to that observed for **147**, indicating that these compounds have similar activities (**Figure 5 – figure supplement 1C**). Three predominant proteins modified by **147-alkyne** in ALMC-2 cells are PDIA1, PDIA4, and PDIA6 (**Figure 5 – figure supplement 1B)** (Paxman et al., 2018). Importantly, addition of excess parent compound **147** competes the labeling of these PDIs by **147-alkyne**, confirming on-target activity (**Figure 5 – figure supplement 1D**) (Paxman et al., 2018). However, we did not observe significant labeling of ALLC in ALMC-2 cells (**Figure 5 – figure supplement 1B,D**) (Paxman et al., 2018). This indicates that **147** influences ALLC secretion from ALMC-2 cells by targeting the activity of PDIs, rather than direct targeting of ALLC. Consistent with this, pharmacologic inhibition of PDIA1 was previously shown to reduce ALLC secretion from HEK293T cells (Cole, Grandjean et al., 2018) and ATF6-dependent reductions in ALLC secretion from HEK293 cells correlate to increased interactions with PDIA4 (Plate et al., 2019). Thus, we sought to define how **147** influences interactions between ALLC and these two PDIs. Interestingly, **147** reduces ALLC interactions with PDIA1 by 25%, while increasing interactions with PDIA4 65% (**Figure 5A,B** and **Figure 5 – figure supplement 1E**). Both of these changes correspond with the alterations in ALLC interactions previously suggested to decrease ALLC secretion (Cole et al., 2018, Plate et al., 2019). This suggests that **147** could reduce ALLC secretion through an ATF6-independent mechanism involving covalent modification of these PDIs.

**Figure 5 (with 1 supplement).**
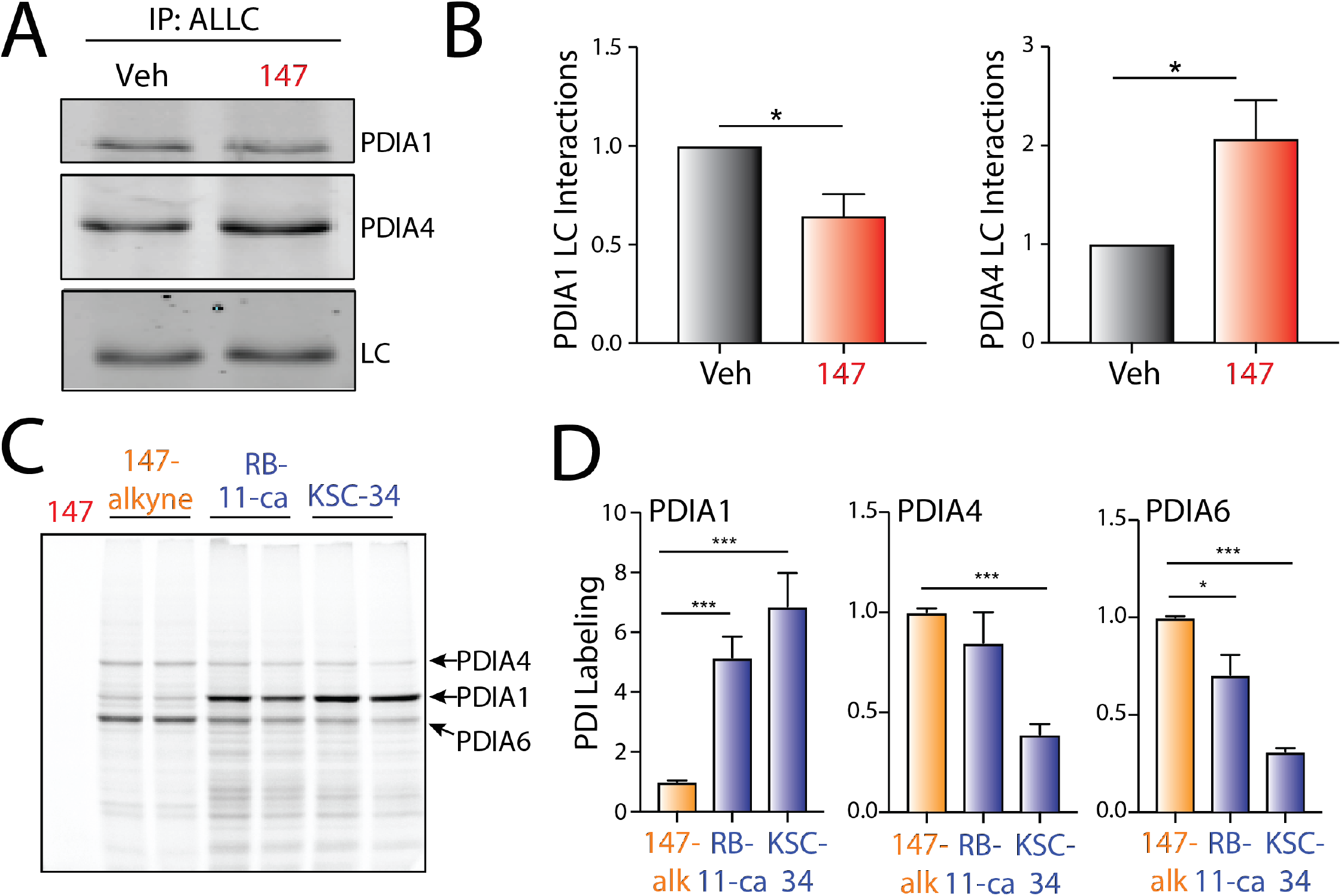
PDI interactions with ALLC are disrupted by 147 treatment. **A.** Representative immunoblot of PDIA1 and PDIA4 in ALLC immunopurifications from ALMC-2 cells treated for 5 hr with vehicle or **147** (10 μM). Inputs from this experiment are shown in **Figure 5 – figure supplement 1E**. **B.** Quantification of PDIA1 and PDIA4 from immunoblots as shown in **Figure 5A**. The recovery of each PDI was normalized to the recovery of ALLC under each condition, allowing accurate evaluation of the interaction between these two proteins. Error bars show SEM for n>5 independent experiments. *p<0.05 for paired t-tests. **C.** Representative fluorescence gel showing the covalent modification of proteins in ALMC-2 cells treated for 18 hr with **147-alkyne** (10 μM), RB-11-ca (30 μM), or KSC-34 (30 μM). Click chemistry was used to incorporate a TAMRA fluorophore onto the alkyne contained in each of these molecules. Cells treated with **147** are shown as a control. The PDI bands were assigned based on previous mass spectrometric analysis of SDS-PAGE bands excised from identical gels of **147-alkyne** treated ALMC-2 cell lysates (Paxman et al., 2018). **D.** Normalized quantification of PDIA1, PDIA4, and PDIA6 labeling from gels as shown in **Figure 5C**. Error bars show SEM for n=4 replicates across two independent experiments. *p< 0.05, **p<0.01, ***p<0.005 for paired t-tests.

RNAi-depletion of PDIs (including *PDIA1* and *PDIA4*) can induce ER stress and subsequent UPR activation (Paxman et al., 2018), challenging our ability to define the specific contributions of individual PDIs in the **147**-dependent reduction in ALLC secretion from ALMC-2 cells. Thus, in order to define how pharmacologic targeting of PDIs influence ALLC secretion, we used structurally distinct PDI inhibitors that covalently modify a similar subset of PDIs to **147**. Recently, the compound RB-11-ca and its more selective analog KSC-34 were identified to covalently modify PDIA1 (**Figure 5 – figure supplement 1F**) (Banerjee, Pace et al., 2013, Cole et al., 2018). These compounds contain an alkyne moiety that allows us to rapidly define their modification of different PDIs using an approach identical to that employed for **147-alkyne** (**Figure 5 – figure supplement 1B**). While RB-11-ca and KSC-34 show increased background labeling relative to **147-alkyne**, both of these compounds label bands that correspond to the three predominant PDIs modified by **147**, PDIA1, PDIA4, and PDIA6, albeit to different extents (**Figure 5C,D**). RB-11-ca and KSC-34 modified PDIA1 to higher levels than **147-alkyne**, consistent with their selectivity for this PDI (Banerjee et al., 2013, Cole et al., 2018). Alternatively, RB-11-ca labeled PDIA4 and PDIA6 to levels 70-85% that observed for **147-alkyne**, whereas KSC-34 showed lower levels of labeling for these PDIs. This is consistent with the increased selectivity of KSC-34 for PDIA1 relative to RB-11-ca (Cole et al., 2018). Collectively, these results show that KSC-34 and, more specifically, RB-11-ca covalently modify similar PDIs to those modified by **147-alkyne**, providing a complementary approach to define how covalent targeting of PDIs impact ALLC secretion from ALMC-2 cells.

### Pharmacologic PDI inhibition reduces ALLC secretion from ALMC-2 cells

Interestingly, both RB-11-ca and KSC-34 reduced ALLC secretion from ALMC-2 cells, as measured by ELISA (**Figure 6A**). RB-11-ca reduced ALLC secretion by 35%, whereas KSC-34 reduced ALLC secretion by 25%. Importantly, this reduction in secretion did not correspond to reductions in ALMC-2 cell viability (**Figure 6 – figure supplement 1A**). Similar reductions in ALLC secretion were observed for RB-11-ca by immunoblotting (**Figure 6 – figure supplement 1B**). As with **147**, RB-11-ca reduced secretion of ALLC dimers and monomers, demonstrating that this PDI inhibitor reduces secretion of ALLC in the conformations most commonly linked to AL pathogenesis (**Figure 6B,C**). This reduction in ALLC secretion also corresponds with reductions in intracellular ALLC measured with immunoblotting (**Figure 6D**) and reductions in protein translation measured by [^35^S] labeling (**Figure 6 – figure supplement 1C**), mimicking the results observed with **147**. Furthermore, RB-11-ca reduced ALLC interactions with PDIA1 and increased interactions with PDIA4 (**Figure 6E,F**). However, neither compound significantly influenced ALLC interactions with PDIA6. Again, these changes are analogous to those observed with **147** (**Figure 5A,B**). However, the reduction in ALLC interactions with PDIA1 was greater than that observed for **147**, likely reflecting the increased labeling of PDIA1 by RB-11-ca (**Figure 5C,D**). These results indicate that pharmacologic targeting PDIs with other PDI inhibitors reduces ALLC secretion from ALMC-2 cells through a mechanism analogous to that observed for **147**. This supports a model whereby **147** reduces ALLC secretion through an on-target ATF6-independent mechanism involving covalent modification of multiple PDIs.

**Figure 6 (with 1 supplement).**
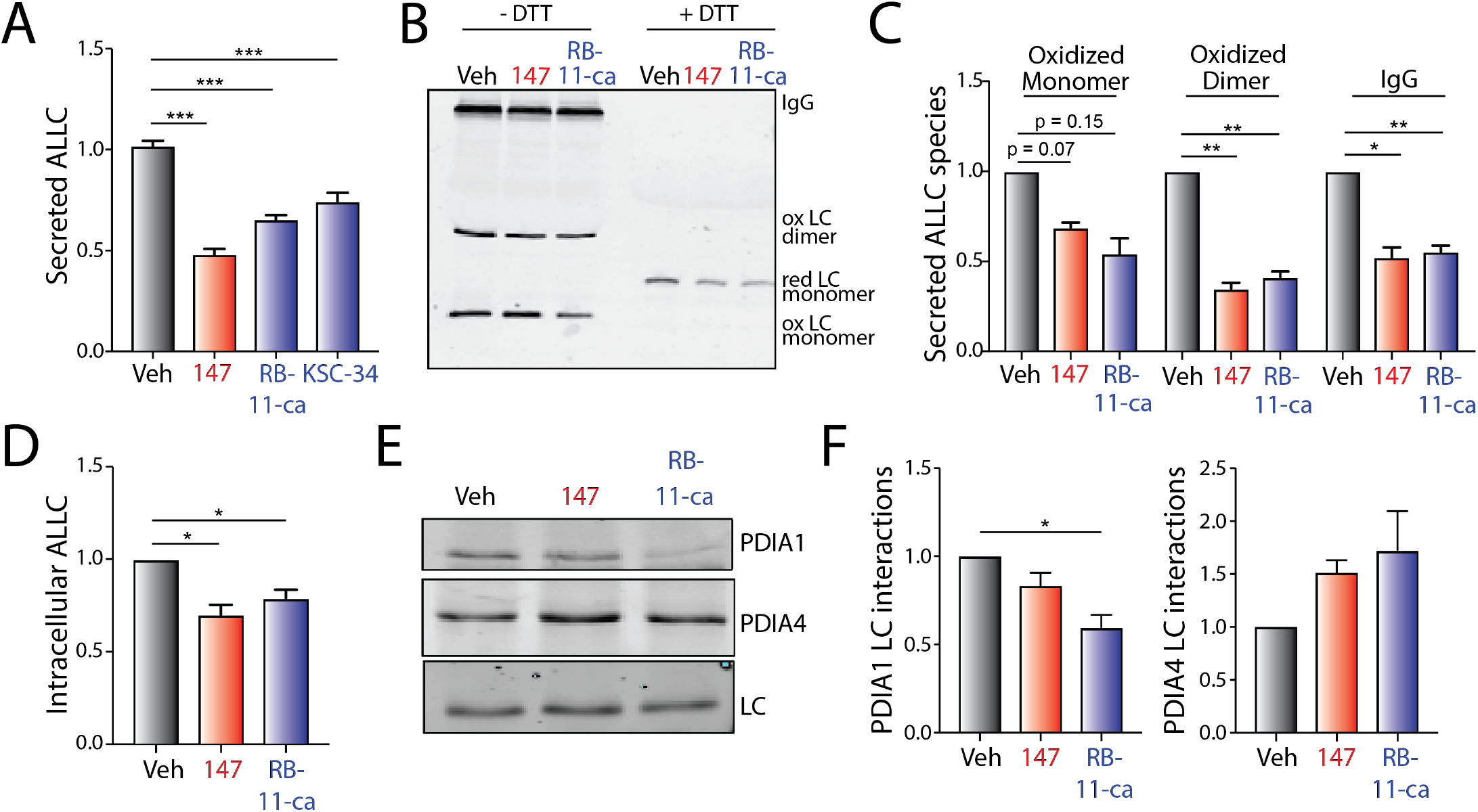
Pharmacologic targeting of PDIs reduces ALLC secretion from ALMC-2 cells. **A.** Graph showing normalized ALLC secretion from ALMC-2 cells treated for 18 hr with vehicle, **147** (10 μM), RB-11-CA (30 μM), or KSC-34 (30 μM). ALLC was quantified by ELISA. Error bars show SEM for n>9 replicates across >two independent experiments. ***p<0.005 from an unpaired t-test. **B.** Representative non-reducing (−DTT) and reducing (+DTT) immunoblots of conditioned media prepared from ALMC-2 cells treated for 18 hr with vehicle, **147** (10 μM) or RB-11-ca (30 μM). The fully-assembled IgGs, oxidized LC dimers, oxidized LC monomers, and reduced LC monomers are indicated. **C.** Graph showing normalized quantification of non-reducing immunoblots as shown in **Figure 6B** for oxidized LC monomers, oxidized LC dimers, and fully-assembled IgGs. Error bars show SEM for n>2 independent experiments. *p<0.05, **p<0.01 from a paired t-test. **D.** Graph showing normalized quantification of ALLC in lysates prepared from ALMC-2 cells treated for 18 hr with vehicle, **147** (10 μM) or RB-11-ca (30 μM). ALLC was measured by immunoblotting. Error bars show SEM for n=4 independent experiments. *p<0.05 from a paired t-test. **E.** Representative immunoblot of ALLC immunopurified from ALMC-2 cells treated for 5 hr with vehicle, **147** (10 μM) or RB-11-ca (30 μM). **F.** Normalized quantification of PDIA1 and PDIA4 from immunoblots as shown in **Figure 6E**. The recovery of each PDI was normalized to the recovery of ALLC under each condition, allowing accurate evaluation of the interaction between these two proteins. Error bars show SEM for n>2 independent experiments. *p< 0.05 from a paired t-test.

### 147-dependent reductions in ALLC secretion are compatible with bortezomib-induced ablation of AL plasma cell

In order to improve the treatment of AL patients in the clinic, any treatment strategy designed to ameliorate AL amyloid pathology must be compatible with standard chemotherapeutics such as the proteasome inhibitor bortezomib used to target the underlying plasma cell malignancy (Gupta, Brauneis et al., 2019, Kastritis et al., 2019, Manwani et al., 2018, Salinaro et al., 2017). Bortezomib induces toxicity in plasma cells through a mechanism involving inhibition of ER-associated degradation and increased ER stress (Nikesitch, Lee et al., 2018, Obeng, Carlson et al., 2006, Ri, 2016). Compound **147** promotes adaptive ER proteostasis remodeling in ALMC-2 plasma cells through both covalent PDI modification and ATF6 activation, suggesting that treatment with this compound could reduce bortezomib-induced toxicity in AL-associated plasma cells. To test this, we evaluated the viability of ALMC-2 cells co-treated with bortezomib and **147**. Bortezomib induces toxicity in ALMC-2 cells with an EC_50_ of 45 nM (**Figure 7A,B**). While **147** treatment alone showed a modest ~15% reduction in ALMC-2 viability (as reported previously (Plate et al., 2016)), this compound did not significantly influence bortezomib-induced toxicity in these cells (**Figure 7A,B**). Furthermore, **147** did not increase or decrease activation of the pro-apoptotic caspases 3/7 in ALMC-2 cells co-treated with subtoxic or toxic doses of bortezomib, respectively (**Figure 7C**). These results indicate that **147** does not interfere with bortezomib-induced plasma cell cytotoxicity and that these two compounds have the potential to be used in combination to mitigate both the amyloid pathology and plasma cell malignancy associated with AL pathogenesis.

**Figure 7.**
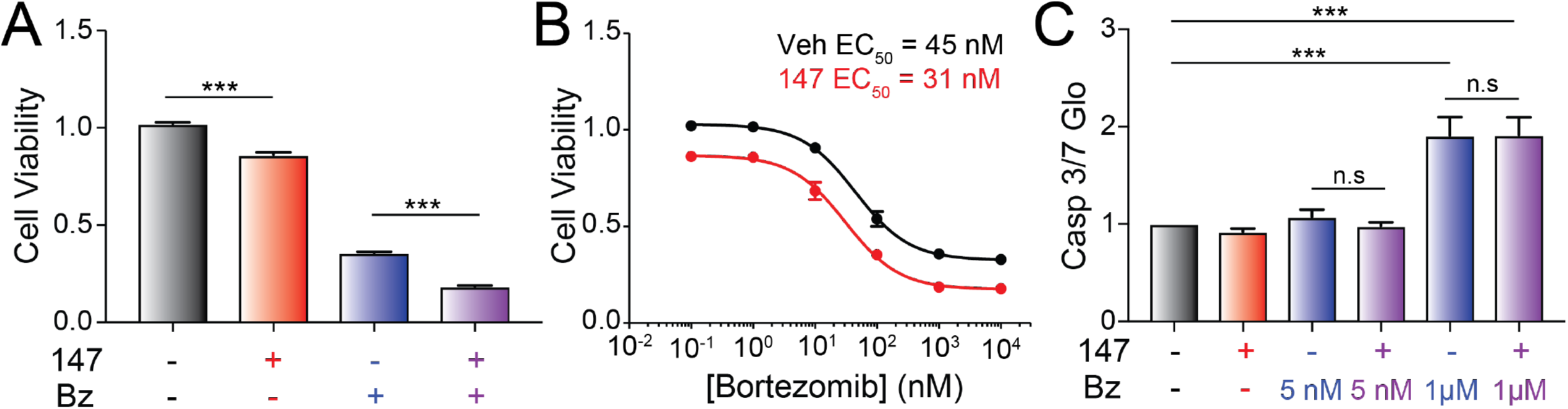
Compound 147 does not influence bortezomib toxicity in ALMC-2 plasma cells. **A.** Graph showing normalized quantification of viability measured by CellTiter-Glo of ALMC-2 cells treated with vehicle, **147** (10 μM) and/or bortezomib (1 μM) for 24h. Error bars show SEM for n=5 replicates. ***p<0.05 from an unpaired t-test. **B.** Graph showing the viability measured by CellTiter-Glo of ALMC-2 cells treated with **147** (10 μM) and/or increasing concentrations of bortezomib (0-10 μM) for 24 h. Error bars show SEM for n=5 replicates. **C.** Graph showing normalized caspase 3/7 activation in ALMC-2 cells treated with **147** (10 μM) and/or the indicated concentration of bortezomib (Bz) for 24 h. Error bars show SEM for n=6 replicates across two independent experiments. ***p<0.005 from an unpaired t-test. (n.s), not significant.

## DISCUSSION

Compound **147** was established as a tool to pharmacologically induce preferential activation of the ATF6 UPR signaling pathway (Paxman et al., 2018, Plate et al., 2016). In this role, **147** has proven useful for probing the therapeutic potential for pharmacologic ATF6 activation in human disease. For example, **147** was used to demonstrate that pharmacologic ATF6 activation protects against ischemia and reperfusion (I/R) injury in both primary cardiomyocytes and in vivo mouse models of myocardial infarction (Blackwood et al., 2019). Genetic depletion or cardiac-specific knockout of *Atf6* blocked **147**-dependent protection in these models, demonstrating that this **147** protection requires ATF6 signaling. Similarly, **147**-dependent ATF6 activation was shown to alter stem cell differentiation, revealing new roles for ATF6 signaling in development (Kroeger, Grimsey et al., 2018). These results highlight the potential for **147**-dependent ATF6 activation to influence physiologic and pathologic processes in diverse biologic contexts.

Here, we show that **147** reduces the plasma cell secretion of destabilized, amyloidogenic ALLC through an on-target, ATF6-*independent* mechanism that involves covalent modification of multiple PDIs. PDIs primarily function in ER proteostasis by regulating the formation of disulfide bonds within secretory proteins, although many PDIs have chaperoning activity that is independent of their redox function (Appenzeller-Herzog & Ellgaard, 2008, Wang et al., 2015). Interestingly, **147** alters the interactions between ALLC and multiple PDIs (e.g., PDIA1 and PDIA4) in ways that are consistent with reduced LC secretion. PDIA1 facilitates LC folding in the early secretory pathway (Borth, Mattanovich et al., 2005, Lilie, McLaughlin et al., 1994). This suggests that reductions in PDIA1 activity should reduce plasma cell secretion of amyloidogenic LCs. Consistent with this, PDIA1 inhibitors decrease ALLC secretion from HEK293 cells (Cole et al., 2018). This indicates that the reduced interaction between ALLC and PDIA1 afforded by **147** contributes to the reduced secretion of ALLC from AL patient plasma cells. Conversely, ALLC increased interactions with PDIA4 in **147**-treated ALMC-2 cells. This is consistent with the increased interactions between ALLC and PDIA4 observed in HEK293 cells following stress-independent ATF6 activation – a treatment where ALLC secretion is reduced through its increased ER retention in chaperone bound complexes (Plate et al., 2019). Interestingly, PDIA4 has been implicated in the retention of misfolded proteins within the ER (Forster, Sivick et al., 2006, Sorensen, Ranheim et al., 2006), suggesting that the increased interaction between ALLC and PDIA4 in **147**-treated ALMC-2 cells likely contributes to the **147**-dependent ER retention of ALLC in ALMC-2 cells. Collectively, these results indicate that **147**-dependent alterations in the activity of multiple PDIs contribute to the reduced secretion of ALLC from AL patient derived plasma cells.

We further defined the importance of PDIs in the plasma cell secretion of ALLC by showing that structurally distinct PDI inhibitors similarly reduce ALLC secretion from ALMC-2 plasma cells. Interestingly, RB-11-ca reduces ALLC secretion through a mechanism analogous to that observed for **147** involving both increased ALLC interactions with PDIA4 and reduced interactions with PDIA1. Furthermore, RB-11-ca reduces intracellular ALLC and induces translational attenuation in ALMC-2 cells, again mirroring results observed for **147**. While the mechanistic basis of this translational attenuation remains to be defined, our results showing that two structurally distinct PDI modifying compounds both reduce protein translation in ALMC-2 cells indicates that this reduced synthesis likely results from PDI modification. Collectively, these results further support a model whereby pharmacologic targeting of multiple PDIs is the primary mechanism responsible for the **147**-dependent reduction in ALLC secretion from ALMC-2 plasma cells.

These results indicate that PDI inhibitors such as **147** or RB-11-ca offer unique opportunities to reduce the secretion and toxic aggregation of amyloidogenic LCs implicated in AL pathogenesis. However, **147** has advantages over other available PDI inhibitors that make it particularly suitable for further translational development. For example, **147** is metabolically activated to its reactive form on the ER membrane (Paxman et al., 2018). This localized activation allows **147** to preferentially modify ER proteins, including many PDIs, minimizing the off-target activity of this compound. This benefit is evident by the reduced background labeling observed when comparing proteins modified by **147** and RB-11-ca. Furthermore, **147** shows reduced labeling of PDIA1 relative to RB-11-ca. Since PDIA1 is important for the folding and trafficking of numerous secretory proteins including insulin (Jang, Pottekat et al., 2019), the reduced targeting of PDIA1 by **147** will minimize potential consequences associated with inhibiting PDIA1 in vivo. Consistent with this, **147** administration has not been associated with any in vivo toxicity to date (Blackwood et al., 2019). Thus, while many different PDI inhibitors have been developed, these unique properties of **147** make it an attractive compound for pharmacologically targeting PDIs to influence ER proteostasis in diseases like AL.

Importantly, we show that **147** reduces secretion of ALLC in dimeric and monomeric conformations – the two conformations most associated with LC proteotoxicity in AL pathogenesis (Andrich et al., 2017, Brenner et al., 2004, Klimtchuk et al., 2010, Ramirez-Alvarado, 2012, Sikkink & Ramirez-Alvarado, 2010). This is consistent with previous results showing that **147** reduces the extracellular aggregation of ALLC secreted from ALMC-2 plasma cells (Plate et al., 2016). Moreover, we demonstrate that **147** does not influence the bortezomib-induced toxicity in ALMC-2 cells, indicating that these two compounds have potential to be used in combination to treat AL in the patients. In this combination approach, **147** would be administered to reduce LC-associated proteotoxicity on distal tissues such as the heart while chemotherapeutics such as bortezomib would be administered to ablate the underlying cancerous plasma cell population. This combined strategy provides the opportunity to both mitigate the LC-associated proteotoxicity and provide chemotherapeutic access to the underlying diseased plasma cells in the 30% of patients suffering from severe LC-associated proteotoxicity. Thus, our results further support the potential application of compounds like **147** for the treatment of AL and reveal new mechanistic insights into compound-dependent reductions in ALLC secretion that will enable the development of next generation compounds with improved translational potential for this disease.

## MATERIALS AND METHODS

### Chemicals

Compound **147**, **147-alkyne**, and all **147** analogs were synthesized in house and are described in (Paxman et al., 2018). RB-11-ca and KSC-34 were both kind gifts from Eranthie Weerapana at Boston College (Banerjee et al., 2013, Cole et al., 2018). Ceapin-7 (CP7) and Ceapin-5 (CP5) were kind gifts from Peter Walter at UCSF (Gallagher et al., 2016, Gallagher & Walter, 2016). Cycloheximide (Fisher), resveratrol (SelleckChem), ISRIB (Sigma), MG132 (Selleckchem), chloroquine (Sigma), PF429242 (S1Pi; Tocris), β-mercaptoethanol (BME; Thermo Fisher), and bortezomib (MilliporeSigma) were all purchased commercially.

### Antibodies

All antibodies were commercially purchased. Antibodies were used for immunoblotting at the indicated dilution: mouse human lambda light chain antibody (Novus Biological, NBP2-29462, 1:250); rabbit PDIA1 (PDI) antibody (GeneTex, GTX101468, 1:1000), rabbit PDIA4 (ERP72) antibody (ProteinTech, 14712-1-AP, 1:1000) and mouse PDIA6 antibody (ProteinTech 66669-1-Ig, 1:1000).

### Cell Lines and Culture Conditions

ALMC-2 and KAS-6/1 plasma cells were a kind gift from Diane Jelinak (Arendt, Ramirez-Alvarado et al., 2008). We cultured these two plasma cell lines in Iscove’s Modified Dulbecco’s Media (IMDM) GlutMAX (Life Technologies) supplemented with penicillin/streptomycin, 5% fetal bovine serum, and 2 ng/mL interleukin-6 (IL-6), following published protocols (Arendt et al., 2008, Plate et al., 2016). Cells were cultured at 37°C with 5% CO_2_.

### ELISA

The ELISA for ALLC and fully-assembled IgGs in conditioned media prepared on ALMC-2 or KAS-6/1 cells was performed using an identical approach to that reported in (Plate et al., 2016). Briefly, ALMC-2 or KAS-6/1 plasma cells were plated 100,000 cells/well in 150 μL of media in 96-well MultiScreen_HTS_ filtration plates (EMD Millipore). Five replicates were treated with DMSO or compounds at the indicated concentrations and incubated for 18 hr. Media was removed by filtration using a QIAvac 96 vacuum manifold (Qiagen) and wells were washed two times with 150 μL media. Wells were then incubated with 150 μL of fresh media for 2 hr and the conditioned media was harvested into a 96-well plate using the vacuum manifold. Whole lysates were obtained by adding RIPA buffer to the cells in the filtration plates. Free LC and IgG concentrations were determined by ELISA in 96-well plates (Immulon 4HBX, Thermo Fisher). Wells were coated overnight at 37 °C with sheep polyclonal free λ LC antibody (Bethyl Laboratories, A80-127A) at a 1:500 dilution or human IgG-heavy and light chain antibody (Bethyl Laboratories, A80-118A) at a 1:2000 dilution in 50 mM sodium carbonate (pH 9.6). In between all incubation steps, the plates were rinsed extensively with Tris-buffered saline containing 0.05% Tween-20 (TBST). Plates were blocked with 5% non-fat dry milk in TBST for 1 hr at 37°C. Media analytes were diluted between 5 – 200 fold in 5% non-fat dry milk in TBST and 100 μL of each sample was added to individual wells. Light chain or IgG standards ranging from 3 – 300 ng/mL were prepared from purified human Bence Jones λ light chain or human reference serum (Bethyl Laboratories, P80-127 and RS10-110). Plates were incubated at 37 °C for 1.5 hr while shaking. Finally, HRP-conjugated goat anti-human λ light chain antibody (Bethyl Laboratories, A80-116P) was added at a 1:5,000 dilution or HRP-conjugate IgG-Fc fragment cross-adsorbed antibody (Bethyl Laboratories, A80-304P, 1:30,000 dilution) was added in 5% non-fat dry milk in TBST, followed by a 1.5 hr incubation of the plates at 37 °C. The detection was carried out with 2,2’-azinobis (3-ethylbenzothiazoline-6-sulfonic acid) (ABTS, 0.18 mg/mL) and 0.03% hydrogen peroxide in 100 mM sodium citrate pH 4.0. Detection solution (100 μL) was added to each well and the plates were incubated at room temperature. The absorbance was recorded at 405 nm and the values for the LC standards were fitted to a 4-parameter logistic function. Light chain or IgG concentrations were averaged from at least 3 independent replicates under each treatment and then normalized to vehicle conditions.

### Immunoblotting for ALLC

Conditioned media was collected from ALMC-2 cells as previously described (Plate et al., 2016). Media was denatured with Laemmli buffer + 100 mM DTT and boiled for 5 min prior to being separated by SDS-PAGE. Conditioned media for non-reducing gels were prepared as above in the absence of DTT. Cell lysates were prepared as previously described (Plate et al., 2016) in RIPA buffer with fresh protease inhibitor cocktail (Roche). Total protein concentration in cellular lysates was normalized using the Bio-Rad protein assay. Lysates were then denatured with Laemmli buffer + 100 mM DTT and boiled for 5 min before being separated by SDS-PAGE. Proteins were then transferred onto nitrocellulose membranes (Bio-Rad) for immunoblotting and blocked with 5% milk in Tris-buffered saline, 0.5 % Tween-20 (TBST). Membranes were then incubated overnight at 4°C with primary antibodies. Membranes were washed in TBST, incubated with IR-Dye conjugated secondary antibodies and analyzed using Odyssey Infrared Imaging System (LI-COR Biosciences). Quantification was carried out with LI-COR Image Studio software.

To measure intracellular ALLC aggregation, ALMC-2 cells were lysed in RIPA buffer as described above. Following centrifugation at 20,000 g for 10 min, we collected the soluble supernatant fraction (SN). We then resuspended the pellet in RIPA + 2% SDS and DTT and sonicated to release aggregate protein. SN and resolubilized pellets were then separated by SDS-PAGE and probed by immunoblotting as above.

### ALLC Immunopurification

ALLC was co-purified with ER proteostasis factors using an identical strategy to that previously reported (Plate et al., 2016). Briefly, ALMC-2 cells were treated for 30 min a room temperature with PBS containing the reversible crosslinker dithiobis (succinimidiyl propionate) (DSP, Thermo Scientific, 500 μM). This crosslinking reaction was quenched by addition of 100 mM Tris pH 7.5 for 15 min. Lysates were then prepared in RIPA buffer, as above. Total protein concentration in cellular lysates was normalized using Bio-Rad protein assay. Cell lysates were then subjected to preclearing with Sepharose 4B beads (Sigma) at 4 °C for 1 h with agitation. Lysates were then incubated overnight at 4 °C with Protein A beads coupled to λ LC antibody, as described below. After four washes in RIPA buffer, crosslinks were cleaved and proteins were eluted by boiling in Laemmli buffer with 100 mM DTT. Eluates were then separated by SDS-PAGE and immunoblotted as described above. Quantifications of proteins co-purified with ALLC were normalized to the recovered ALLC, as described previously (Plate et al., 2019).

For immunopurification, protein A Sepharose 4B beads (101041, Thermo Fisher) were coupled to human lambda light chain antibody (A80-112A, Bethyl laboratories) (Plate et al., 2016). Briefly, beads were washed twice with RIPA buffer (50 mM Tris, pH 7.5, 150 mM NaCl, 0.1 % SDS, 1% Triton X-100, 0.5% deoxycholate) and resuspended in 1 mL of PBS. Beads were then incubated with primary rabbit human lambda light chain antibody (40 uL of antibody for 70-90-μL of beads) and rocked for 1 hour at room temperature. Beads were washed 3 times with 0.2M borate buffer pH = 9. The antibody was crosslinked to the beads by incubating with 20 mM of the crosslinker dimethyl pimelimidate (DMP, Thermo Scientific, 20 μM) for 30 min at room temperature. The crosslinking reaction was stopped by incubation with 0.2 M ethanolamine pH 8 for 2 h at room temperature. Beads were then washed with Glycine buffer (0.1M, pH=3) to remove uncrosslinked antibody, followed by two washes with PBS. Finally, beads were resuspended in storage buffer (PBS, Azide (0.01%)) and stored at 4°C.

### Quantitative RT-PCR

The mRNA levels of target genes were measured using quantitative RT-PCR on an ABI 7900HT Fast Real Time PCR machine using the same primers and identical approach to that published in (Plate et al., 2016).

### Cell Viability and Apoptosis Assays

ALMC-2 cells were plated at 33,000 cells/well in a translucent, flat-bottomed 96 well plate, and treated for 18 hr with vehicle, 147 (10 μM) or bortezomib (at the indicated doses). After incubation, cell metabolic activity or caspase 3/7 activity was measured using the CellTiter-Glo assay (Promega) and Caspase-Glo 3/7 assay (Promega), respectively, according to the manufacturer’s instructions. Briefly, the plates containing cells were removed from the incubator and allowed to equilibrate to room temperature for 30 min. CellTiter-Glo or Caspase-Glo reagent was added to each well at a 1:1 v/v ratio and incubated for 2 min on an orbital shaker to induce cell lysis. The plate was then incubated at room temperature for 10 min to stabilize the luminescent signal and read on a Tecan F200 Pro microplate reader.

### [^35^S] Metabolic Labeling

For metabolic labeling experiments, ALMC-2 cells were plated at a density of 1×10^6^ cells/well on a 6 well plate. Cells were then treated with **147** (10 μM), RB-11-ca (30 μM), and/or resveratrol (10 μM), as indicated. Cells were transferred to microfuge tubes, washed with PBS, and metabolically labeled in DMEM-Cys/-Met (Corning) supplemented with glutamine, penicillin/streptomycin, dialyzed fetal bovine serum, and EasyTag EXPRESS [^35^S] Protein Labeling Mix (Perkin Elmer) for 30 min in a tissue culture incubator. Cells were washed twice with PBS and lysates were harvested using RIPA buffer (50 mM Tris, pH 7.5, 150 mM NaCl, 0.1 % SDS, 1% Triton X-100, 0.5% deoxycholate) and protease inhibitor cocktail (Roche). ALLC was immunopurified using human lambda LC coupled to Protein A Sepharose beads as described above. Lysates were incubated with the beads overnight at 4 degrees. Beads were washed 5 times with RIPA buffer. ALLC was then isolated by boiling in Laemmli buffer + 100 mM DTT and separated on SDS-PAGE. For whole cell lysates, cells were lysed in RIPA buffer. Laemmli buffer + 100 mM DTT was then added to the lysates and separated by SDS-PAGE. All gels were dried, exposed to phosphorimager plates (GE Healthcare), and imaged with a Typhoon imager. Band intensities were quantified by densitometry in ImageQuant.

### Profiling of Targets for 147 and PDI Inhibitors

Proteins labeled by treatment with **147-alkyne**, RB-11-ca, or KSC34 in were performed as previously described (Paxman et al., 2018). Briefly, ALMC-2 cells in 6-well plates were treated for 18 h with **147-alkyne** (10uM), RB-11-ca (30 μM) or KSC-34 (30 μM). Lysates were then prepared in RIPA buffer as described above. Protein concentrations were then quantified by Bradford. Lysates containing 50 μg of protein were then incubated for 2 h with shaking at 37°C with 100 μM TAMRA azide (Click Chemistry Tools Scottsdale, Az), 800 μM copper(II) sulfate, 1.6 mM BTTAA ligand (2-(4-((bis((1-tert-butyl-1H-1,2,3-triazol-4-yl)methyl)amino)methyl)−1 H-1,2,3-triazol-1-yl)acetic acid) (Click Chemistry Tools Scottsdale, Az), and 5 mM sodium ascorbate. Samples were then boiled with Laemmli Buffer and 100mM DTT and separated by SDS-PAGE. TAMRA-labeled proteins were then imaged using a Bio-Rad ChemiDoc using the rhodamine channel.

### Statistical Methods

Replicates for each experiment comprise cells in separate wells/dishes that were treated independently. The number of replicates and independent experiments for each figure panel are clearly stated in the figure legends. Unless otherwise noted, all p-values were calculated by performing a two-tailed paired or two-tailed unpaired t-test.

## ACKNOWLEDGEMENTS

We thank Evan Powers and Jessica Rosarda for critical reading of the manuscript and Kelly Chen for experimental support. We thank Diane Jelinak (Mayo) for providing the ALMC-2 and KAS-6/1 cell lines, Eranthie Weerapana (Boston College) for RB-11-ca and KSC-34, and Peter Walter (UCSF) for providing CP7 and CP5. This work was funded by the NIH (DK107604 to RLW), a Leukemia & Lymphoma Society Postdoctoral Fellowship (to BR), and an American Heart Association Predoctoral Fellowship (to ICR).

## AUTHOR CONTRIBUTIONS

BR, JSM, ICR, and RLW conceptualized the experiments and performed formal analysis of the primary data. BR, JSM performed the experiments. RJP and JWK provided resources and developed methodology used in this manuscript. BR and RLW wrote the manuscript and all authors were involved in the review and editing. RLW was responsible for supervision, administration, and funding acquisition for the work described in this manuscript.

---

**Figure 1 – figure supplement 1.**
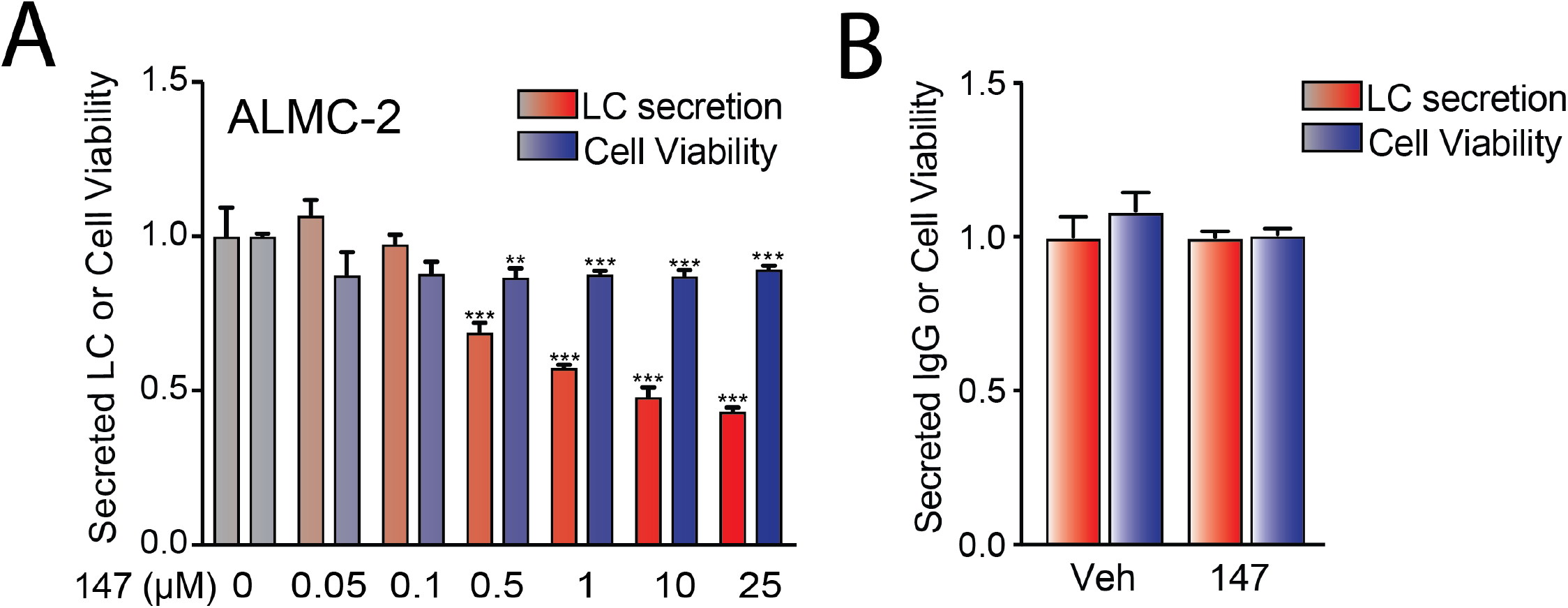
Compound 147 reduces secretion of ALLC from AL patient derived ALMC-2 cells. **A.** Graphs showing relative ALLC in conditioned media measured by ELISA (red) and cellular viability measured by CellTiter-Glo (blue) in ALMC-2 cells treated for 18 hr with vehicle or **147** (10 μM). Error bars show SEM for n= 4 replicates. ***p<0.005 vs Veh from an unpaired t-test. **B.** Graph showing relative media levels of IgG measured by ELISA (red) and cellular viability measured by CellTiter-Glo (blue) in KAS-6/1 cells treated with vehicle or **147** (10 μM) for 18 hr. Error bars show SEM for n = 5 replicates. ***p<0.005 vs Veh from an unpaired t-test.

**Figure 2 – figure supplement 1.**
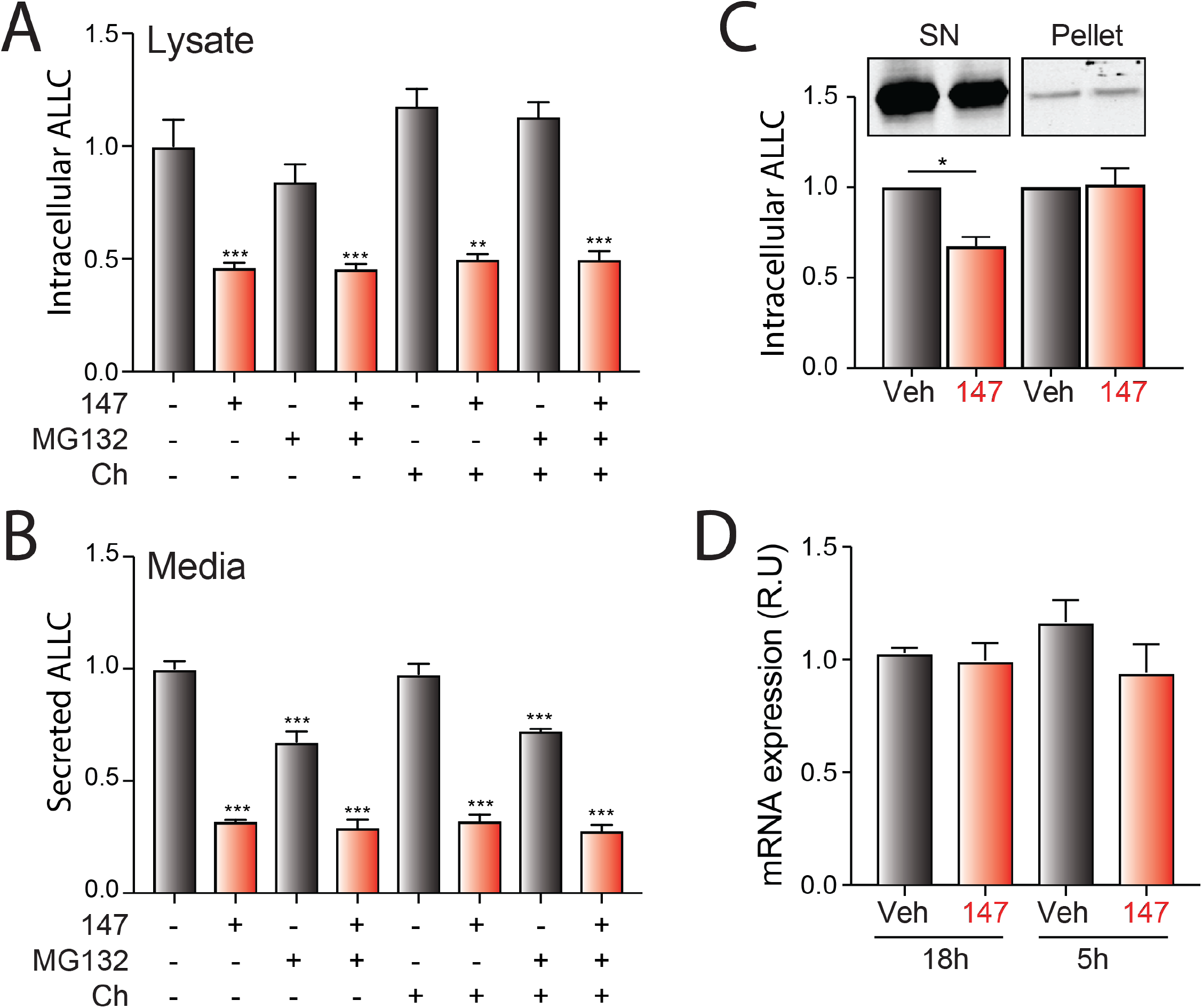
Compound 147 reduces lysate levels of ALLC in ALMC-2 cells. **A.** Bar graphs showing normalized amounts of ALLC in lysates prepared from ALMC-2 cells treated for 18 hr with vehicle or **147** (10 μM). MG132 (10 μM) or chloroquine (Ch, 20 μM) were added as indicated for the last 5 hours of this incubation and during the 2 hours of media conditioning. ALLC was quantified by ELISA. Error bars show SEM for n=5 replicates. *** p<0.005 vs Veh from an unpaired t-test. **B.** Bar graphs showing normalized amounts of ALLC in conditioned media prepared from ALMC-2 cells treated for 18 hr with Veh, **147** (10 μM). MG132 (10 μM) or chloroquine (20 μM) were added for the last 5 hours of this incubation and during the 2 hours of media conditioning. ALLC was quantified by ELISA. Error bars show SEM for n=5 replicates. *** p<0.005 vs Veh from an unpaired t-test. **C.** Representative immunoblot and normalized quantification of ALLC in fractionated lysates from ALMC-2 cells treated with Veh or **147** (10 μM) for 18 hr. The supernatant fraction (SN) is the soluble protein fraction obtained after lysing with RIPA buffer (no DTT). Pellets derived from the first lysis were then solubilized by treatment with SDS (2%), DTT and sonication. Error bars show SEM for n=3 independent replicates. *p<0.05 from a paired t-test **D.** Bar graphs showing ALLC mRNA measured by qPCR in ALMC-2 cells treated with vehicle or **147** for 18 hr or 5 hr. Error bars show 95% confident intervals for n=3 replicates.

**Figure 3 – figure supplement 1.**
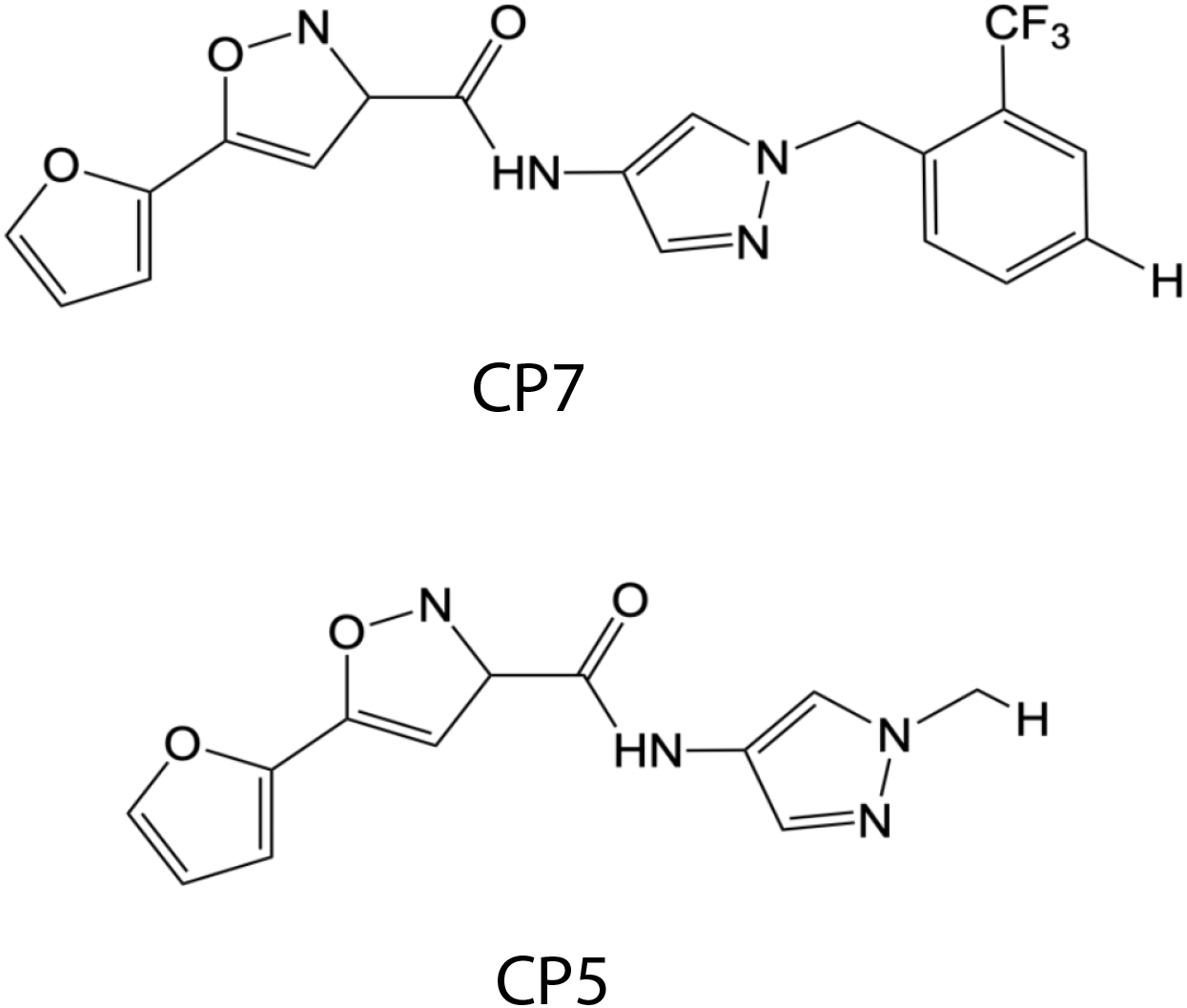
147-Dependent Reductions in ALLC Secretion are Independent of ATF6 Activation. Chemical structures of the ATF6 inhibitor CP7 and its inactive analog CP5 (Gallagher et al., 2016, Gallagher & Walter, 2016).

**Figure 4 – figure supplement 1.**
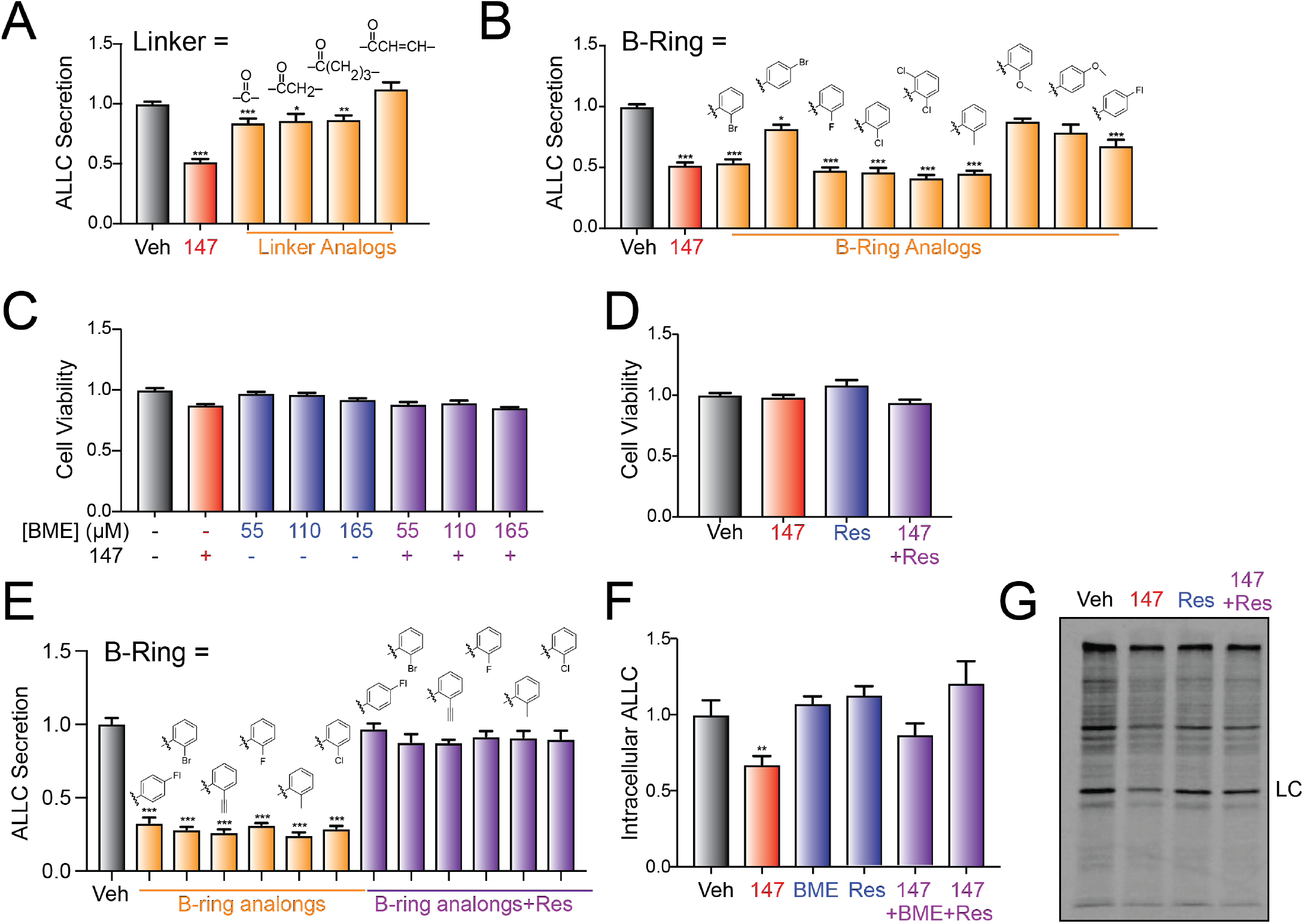
Compound 147 reduces ALLC secretion through a mechanism involving compound metabolic activation and covalent protein modification. **A.** Graph showing normalized ALLC in conditioned media from ALMC-2 cells treated for 18 hr with vehicle, **147** (10 μM) or the indicated **147** ‘Linker’-analog (10 μM). ALLC was quantified by ELISA. Error bars show SEM for n=10 replicates across two independent experiments. *p< 0.05, **p<0.01, ***p<0.005 vs Veh from an unpaired t-test. **B.** Graph showing normalized ALLC in conditioned media from ALMC-2 cells treated for 18 hr with vehicle, **147** (10 μM) or the indicated **147** ‘B’ ring’-analog (10 μM). ALLC was quantified by ELISA. Error bars show SEM for n=10 replicates across two independent experiments. *p< 0.05, **p<0.01, ***p<0.005 vs Veh from an unpaired t-test. **C.** Graph showing normalized quantification of cell viability measured with CellTiter-Glo of ALMC-2 cells treated with vehicle, **147** (10 μM) and/or β-mercaptoethanol (BME) for 18 hr. Error bars show SEM for n=5 replicates.. **D.** Graph showing normalized quantification of cell viability measured with CellTiter-Glo of ALMC-2 cells treated with vehicle, **147** (10 μM) and/or resveratrol (Res: 10 μM) for 18 hr. Error bars show SEM for n=10 replicates across two independent experiments. **E.** Graph showing normalized ALLC in conditioned media from ALMC-2 cells treated for 18 hr with vehicle, **147** (10 μM) or ‘B ring’-analog (10 μM) with or without resveratrol (10 μM). Error bars show SEM for n>3 replicates. ***p<0.005 vs Veh from an unpaired t-test. **F.** Graph showing the normalized amounts of ALLC in lysates prepared from ALMC-2 cells treated for 18 hr with vehicle, **147** (10 μM), resveratrol (10 μM), or β-mercaptoethanol (BME: 165 μM), as indicated. ALLC was quantified by ELISA. Error bars show SEM shows n=5 replicates. **p<0.01 vs Veh from an unpaired t-test. **G.** Representative autoradiogram of whole cell lysates prepared from ALMC-2 cells treated for 18 hr with vehicle, **147** and/or resveratrol (10 μM) and then metabolically labeled with [^35^S] for 30 min.

**Figure 5 – figure supplement 1.**
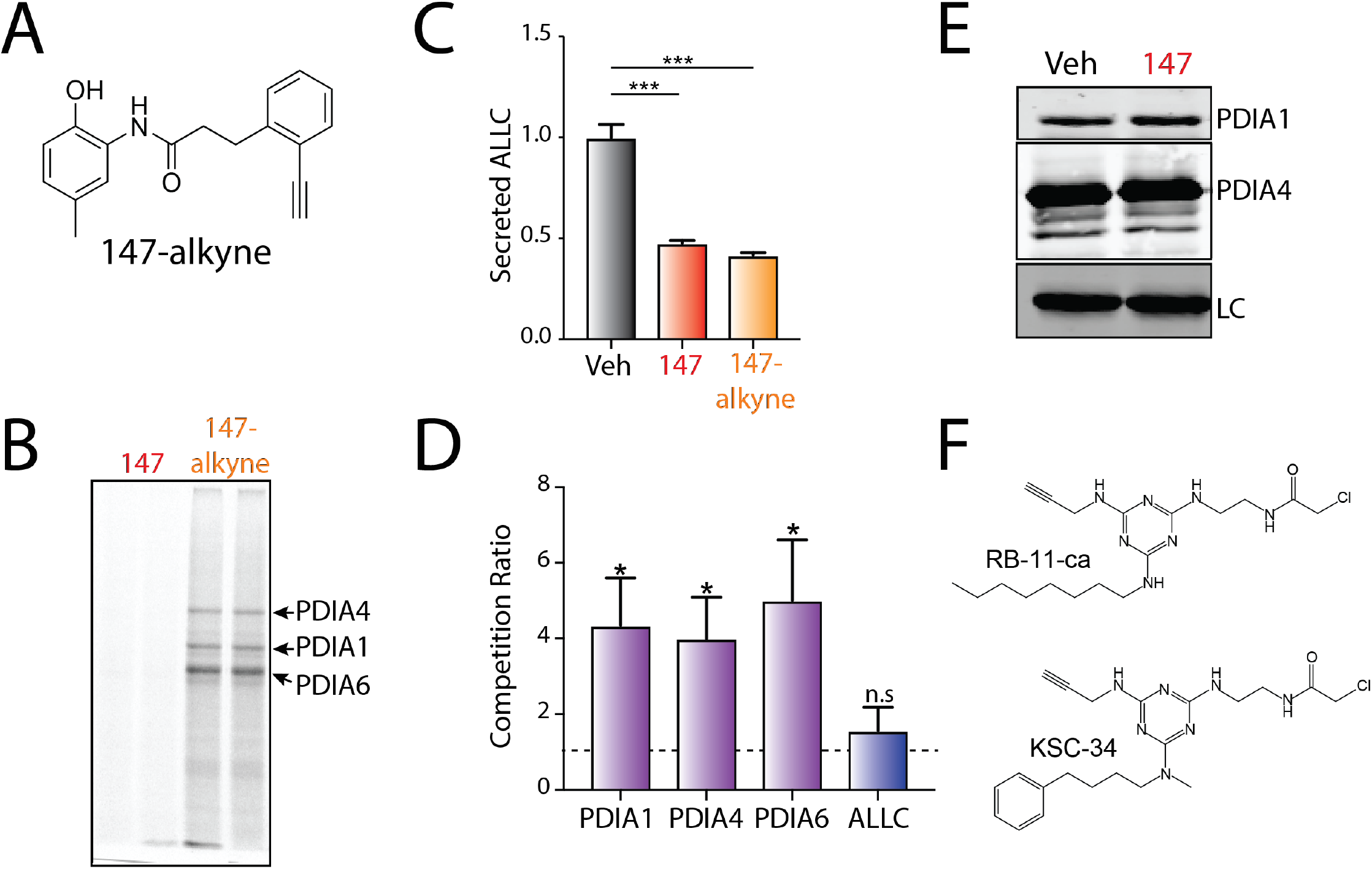
PDI interactions with ALLC are disrupted by 147 treatment. **A.** Chemical structure of **147-alkyne**. **B.** Representative gel showing the covalent modification of proteins in ALMC-2 cells treated for 18 hr with **147** (10 μM) or **147-alkyne** (10 μM). Click chemistry was used to incorporate a TAMRA fluorophore onto the alkyne contained in each of these molecules. Cells treated with **147** are shown as a control. The PDI bands were assigned based on previous mass spectrometric analysis of SDS-PAGE bands excised from identical gels of **147-alkyne** treated ALMC-2 cell lysates (Paxman et al., 2018). **C.** Graph showing ALLC secreted from ALMC-2 cells treated for 18 hr with vehicle, **147** (10 μM) or **147**-alkyne (10 μM). Error bars show SEM for n=10 replicates across two independent experiments. ***p<0.005 from an unpaired t-test. **D.** Graph showing competition ratio of the indicated PDI or ALLC in affinity purifications isolated from ALMC-2 cells treated for 18 h with **147-alkyne** (10 μM) or **147-alkyne** (10 μM) and an excess of **147** (50 μM). These data are from experiments described in (Paxman et al., 2018). Briefly, after incubation with ALMC-2 cells, **147-alkyne** was modified with a biotin allowing isolation of the proteins covalently modified by **147-alkyne** using streptavidin affinity purification. The recovery of different proteins across conditions was then quantified using tandem mass tag (TMT) labeling and multi-dimensional protein identification technology (MuDPIT). The competition ratio was calculated as previously described using the equation: competition ratio = protein signal from cells treated with **147-alkyne** treated / protein signal from cells treated with **147-alkyne** and excess **147** (Paxman et al., 2018). Error bars show SEM for n=4 paired replicates across two independent experiments. *p<0.05 from a paired t-test. (n.s), not significant. **E.** Immunoblot showing lysate levels of PDIA1, PDIA4 and ALLC in ALMC-2 cells treated for 5 hr with vehicle or **147** (10 μM). These are inputs from the immunopurifications shown in **Figure 5A**. **F.** Chemical structures of the PDI inhibitors RB-11-ca and KSC-34.

**Figure 6 – figure supplement 1.**
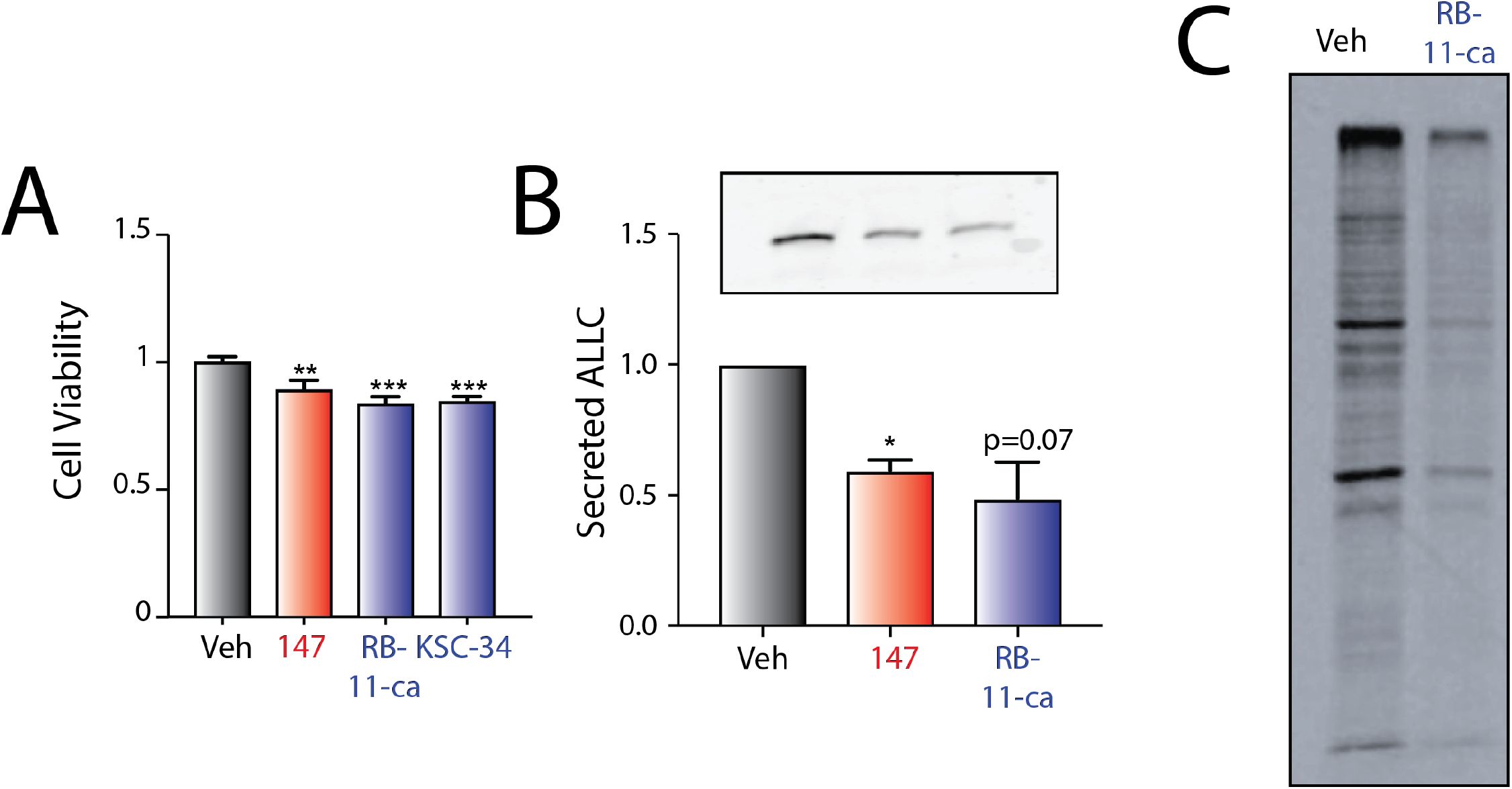
Pharmacologic targeting of PDIs reduces ALLC secretion from ALMC-2 cells. **A.** Graph showing the viability measured by CellTiter Glo for ALMC-2 cells treated for 18 hr with vehicle, **147** (10 μM), RB-11-ca (30 μM), or KSC-34 (30 μM). ALLC was quantified by ELISA. Error bars show SEM for n=11 replicates across 2 independent experiments. ***p<0.005 vs Veh from an unpaired t-test. **B.** Representative immunoblot and quantification of ALLC in conditioned media prepared from ALMC-2 cells treated with vehicle, **147** (10 μM) or Rb-11-ca (30 μM) for 18 h. Error bars show SEM for n=3 independent experiments. *p<0.05 vs Veh from a one-tailed paired t-test. **C.** Representative autoradiogram of whole cell lysates prepared from ALMC-2 cells treated for 18 h with vehicle, **147** (10 μM) or Rb-11-ca (30 μM) and then metabolically labeled with [^35^S] for 30 min.

